# TMEM63 proteins act as mechanically activated cholesterol modulated lipid scramblases contributing to membrane mechano-resilience

**DOI:** 10.1101/2025.07.26.664997

**Authors:** Yiechang Lin, Zijing Zhou, Yaoyao Han, Delfine Cheng, Haoqing Wang, Arnold Lining Ju, Yixiao Zhang, Charles D Cox, Ben Corry

**Affiliations:** Research School of Biology, Australian National University, Acton, ACT 2601, Australia; Victor Chang Cardiac Research Institute, Sydney, NSW, 2010, Australia; School of Clinical Medicine, Faculty of Medicine and Health, University of New South Wales, Sydney, NSW 2052, Australia; Interdisciplinary Research Center on Biology and Chemistry, Shanghai Institute of Organic Chemistry, Chinese Academy of Sciences, China; State Key Laboratory of Chemical Biology, Shanghai Institute of Organic Chemistry, Chinese Academy of Sciences, Shanghai, 200032, China; School of Biomedical Engineering, The University of Sydney, Darlington, NSW, 2008, Australia; School of Biomedical Sciences, Faculty of Medicine & Health, UNSW Sydney, Kensington, NSW, 2052, Australia

**Author notes:** Corresponding authors: & &. These authors contributed equally (Yiechang Lin, Zijing Zhou, Yaoyao Han).

## Abstract

OSCA/TMEM63 mechanosensitive ion channels play critical physiological roles in plants and animals. These channels bear structural homology to the dual functional TMEM16 family, and OSCA1.2 was recently shown to form a lipid-lined ion conduction pathway in the open state. This raised the question of whether members of the OSCA/TMEM63 family may also function as mechanically activated lipid scramblases. Using a combination of in vitro and cellular assays with computational techniques, we show that phospholipids can be translocated through the open pores of OSCA1.1/1.2/2.2 and TMEM63A/B proteins, suggesting a dual ion channel and lipid scramblase function for members of this protein family. We characterize the effects of mutating key groove lining residues demonstrating that different residues form bottlenecks for lipids and ions respectively and show that cholesterol inhibits lipid scrambling by stabilizing the closed state and slowing translocation through the open pore. We show that lipid scrambling in TMEM63 proteins can be activated by mechanical forces in the membrane, making these the first identified mechanically activated lipid scramblases. Finally, we demonstrate that this activity is important for the mechanically induced morphological remodeling of biological membranes and the resilience of cells to high mechanical forces.

## Introduction

Mechanosensitive ion channels are ubiquitously expressed across all domains of life and play essential roles in bacteria, plants and animals^1^. Proteins belonging to the OSCA/TMEM63 family form the largest known group of mechanosensitive (MS) channels^2,3^. Like other structurally unrelated MS channels, OSCA/TMEM63 are activated by membrane tension^2,4,5^, according to the force-from-lipids principle^6^. In plants, OSCA channels are essential for the osmotic stress response^7–9^, with more than 15 paralogues identified in *Arabidopsis thaliana*^2,3,7^. In animals, the homologous TMEM63 proteins are reported to be involved in myelination^10^, hearing^11^, detection of food grittiness^12^ and the thirst response^13^. Despite the structural similarity, OSCA channels adopt a dimeric assembly with each monomer containing an ion conducting pore^5,14–17^, whereas TMEM63 channels function as monomers^4,18,19^. While most of the 15 OSCA paralogues display stretch activated currents with a similar threshold for activation, they have considerable diversity in their single-channel conductance ranging from >100pS for OSCA1.1 and 1.2 to below detectable limits for OSCA2.3^2^. TMEM63 proteins are reported to have higher thresholds for mechanical activation, and single channel conductance below detectable limits^2,4^.

OSCA and TMEM63 channels share structural homology to the TMEM16 family of calcium activated chloride channels and phospholipid scramblases. TMEM16 scramblases mediate the ATP-independent flipping of phospholipids between membrane leaflets^20–23^. This leads to externalization of phosphatidylserine^24^, usually found only in the intracellular leaflet, contributing to processes such as blood coagulation in platelets, bone development and cell fusion^21,25–27^. Computational studies support a model where lipid translocation in TMEM16 scramblases occurs via the ‘credit card’ mechanism, in which charged lipid headgroups move through a membrane-spanning hydrophilic groove of the protein in the open state^23,28,29^. Consistent with the dual ion channel and scramblase activity of some TMEM16 proteins, the translocating phospholipid headgroups are thought to interact with groove residues to form a proteo-lipidic pore which facilitates the simultaneous permeation of ions^21,22,30^. Recent work indicates that OSCA channels possess a lipid-lined ion conduction pathway in which 6 to 7 lipids form a belt connecting the two membrane leaflets^5^. Similarly, a lipid-lined pore has been proposed for another related channel family - the TMC proteins.^31,32^ This, combined with the structural homology between OSCA/TMEM63, TMEM16 and TMC proteins, raises the question of whether OSCA/TMEM63 channels can also facilitate lipid translocation between membrane leaflets upon mechanical force. While previous work has suggested scrambling only in TMEM63B^33^, only in mutant TMEM63 proteins^34,35^ or in response to Ca^2+^ ^36^ there is no evidence for the native stimuli for TMEM63 lipid scrambling or if lipid scrambling by these proteins plays a physiological role.

To assess the ability of OSCA/TMEM63 proteins to scramble lipids as well as the activating stimuli and physiological role of lipid scrambling, we combined molecular dynamics simulations of cryo-EM and AlphaFold2 predicted structures of these channels, *in vivo* and *in vitro* scramblase assays, together with structure determination using cryo-electron microscopy (cryo-EM). We demonstrate that OSCA1.1/1.2/2.2 and TMEM63A/B can translocate lipids, that this can be inhibited by cholesterol, and that most importantly lipid scrambling can be activated in wild-type proteins by mechanical forces driving morphological changes in model membranes. At native protein levels this scrambling can then protect membranes from high mechanicals force indicative of a role in membrane mechano-resilience.

## Results

### OSCA1.2 and TMEM63A can scramble lipids

To assess the ability of OSCA1.2 to scramble lipids, we carried out 20 μs long coarse-grained molecular dynamics (CG-MD) simulations of OSCA1.2 in its closed and open conformations as well as the open structure of nhTMEM16, a bona fide scramblase. During simulations of the open state, lipids spontaneously form a belt linking the upper and lower membrane leaflets as seen in recent all-atom simulations, with a water filled pore forming between the lipids and protein (Fig. 1A). While no lipid translocation events occur across the closed OSCA1.2, these happen spontaneously in simulations of the open structure of OSCA1.2 at a rate greater than for nhTMEM16 (Fig. 1B, Supplementary Video 1).

**Figure 1.**
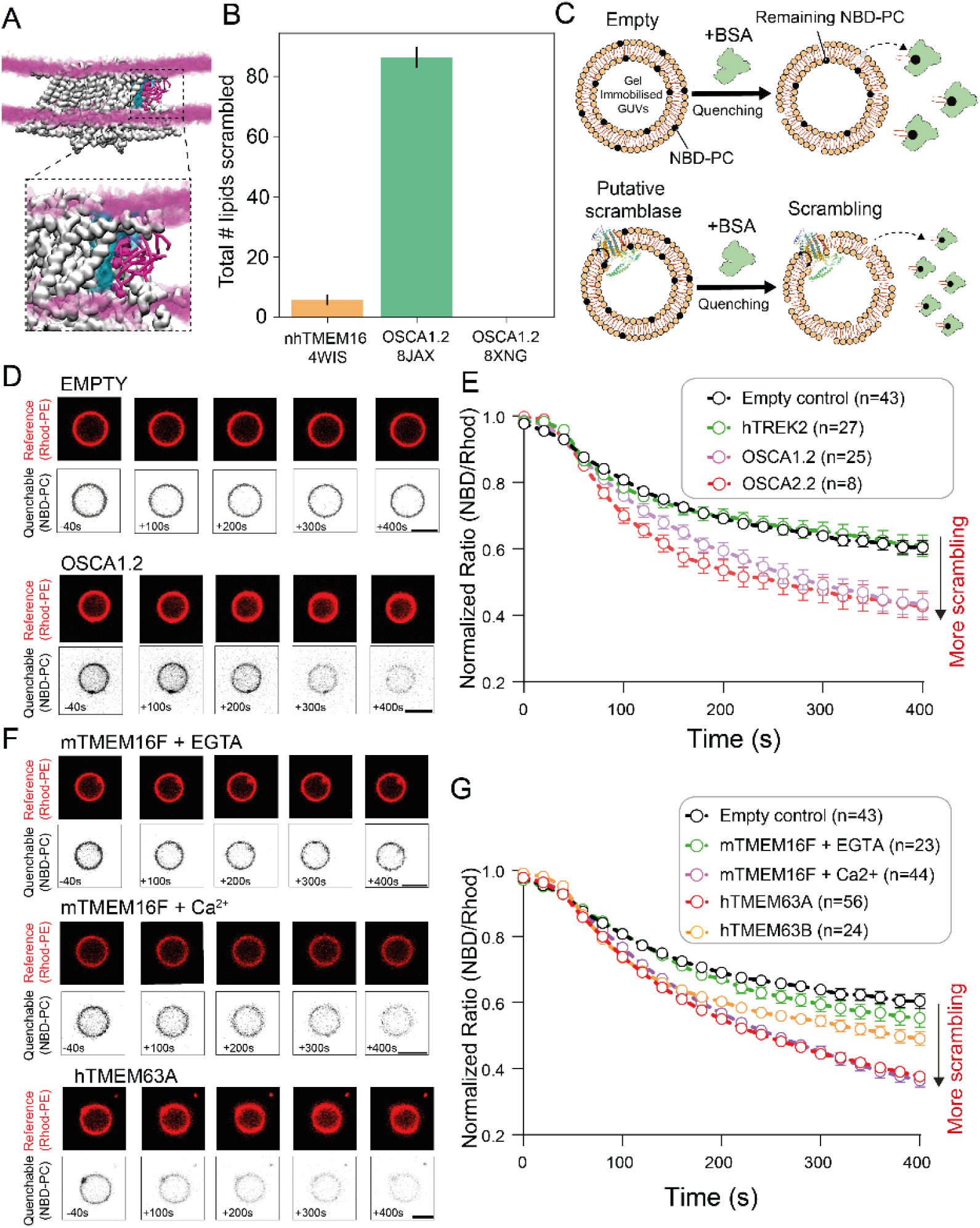
OSCA1.2/2/2 and TMEM63A/B can scramble lipids. (A) CG-MD simulations of OSCA1.2 show a lipid lined and water filled pore. Protein shown in white, lipid headgroups in the bilayer by the purple surface, lipids forming the pore wall as purple licorice, and the water filled pore in blue. (B) Total number of lipids scrambled in 20 µs CG-MD simulations for closed (PDB ID: 8XNG), and open OSCA1.2 (PDB ID:8XAJ) as well as open nhTMEM16 (PDB ID: 4WIS). Error bars represent SEM (n=3). (C) Schematic of *in vitro* scramblase assay using giant unilamellar vesicles (GUVs) and bovine serum albumin (BSA) induced quenching to measure putative scramblase activity. (D) Confocal images of soy polar lipid GUVs containing 0.2 % w/w Rhod-PE (1,2-dioleoyl-sn-glycero-3-phosphoethanolamine-N-(lissamine rhodamine B sulfonyl) and 0.8 % w/w NBD-PC (1-palmitoyl-2-{6-[(7-nitro-2-1,3-benzoxadiazol-4-yl)amino]hexanoyl}-*sn*-glycero-3-phosphocholine) before and after addition of 4 mg/mL BSA (scale bar = 5 µm). (E) Time course of changes in normalized ratio of NBD/Rhod fluorescence intensity after BSA addition in empty GUVs (n=43) and those reconstituted with hTREK2 (n=18), OSC1.2 and OSCA2.2 Data points represent mean ± SEM. (F) Confocal images of soy polar lipid GUVs containing 0.2 % w/w Rhod-PE (1,2-dioleoyl-sn-glycero-3-phosphoethanolamine-N-(lissamine rhodamine B sulfonyl) and 0.8 % w/w NBD-PC (1-palmitoyl-2-{6-[(7-nitro-2-1,3-benzoxadiazol-4-yl)amino]hexanoyl}-*sn*-glycero-3-phosphocholine) before and after addition of 4 mg/mL BSA (scale bar = 5 µm). (G) Time course of changes in normalized ratio of NBD/Rhod fluorescence intensity after BSA addition in empty GUVs (n=43) and those reconstituted with mTMEM16F with 2 mM EGTA (n=23), mTMEM16F with 1 mM Ca^2+^ (n=44), hTMEM63A (n=46), hTMEM63B (n=24).

Given the *in silico* results show the potential for lipid scrambling in OSCA1.2, we next developed an *in vitro* assay to measure lipid scrambling. We generated giant unilamellar vesicles (GUV) containing a fraction of fluorescent lipids with both tail labelled nitrobenzoxadiazole (NBD) and head labelled Rhodamine B (Rhod). The change in GUV fluorescence ratio of NBD/Rhod was then measured before and after the addition of bovine serum albumen (BSA) that only quenches NBD exposed on the outer leaflet (Fig.1C). In the absence of lipid scrambling, fluorescence is expected to drop to close to half its initial value after the addition of BSA, whereas it will decline further if lipids are moving from the inner to outer leaflet. To record these individual GUVs over time we developed a new method of agarose gel-based GUV immobilization using low melting point agarose. Our results show that NBD fluorescence declined significantly more in OSCA1.2 and OSCA2.2 containing liposomes than in empty controls and GUVs containing hTREK2 (a mechanically activated ion channel known not to scramble lipids). (Fig.1D-E). This is indicative of OSCAs scrambling lipids. We also characterize the ion conductance of OSCA2.2 and its electrophysiological characteristics, including its small peak currents and lack of desensitization, look much more similar to TMEM63 proteins than to OSCA1.1/1.2 (Fig S1). We saw even more robust evidence of OSCA1.2/1.1 lipid scrambling when reconstituted into GUVs composed of pure DOPE/POPC at a ratio of 7:3, again compared to rigorous positive and negative controls (Fig.S2A-B). Interestingly, GUVs containing hTMEM63A or hTMEM63B also showed a reduction in fluorescence ratio significantly below the negative controls. (Fig.1F-G). As the best control we used GUVs containing the known Ca^2+^ activated scramblase mTMEM16F. We saw no scrambling in the presence of 2 mM EGTA (Fig.1F-G) but a significant drop in NDB/Rhod fluorescence, similar to that seen for TMEM63 proteins, for mTMEM16F in the presence of 1 mM Ca^2+^. This provides strong evidence from a reductionist system that lipid scrambling may be a common feature of wild-type proteins in this family.

### Predicted open structures of hTMEM63A scramble lipids and have a lipid lined pore which conducts ions

To find potential structures of alternate conformations of the OSCA/TMEM63 family, we generated 100 structures each of OSCA1.2, hTMEM63A, hTMEM63B and hTMEM63C using AF2. Remarkably, AF2 can sample multiple conformational states of these proteins without the need for MSA subsampling nor templating to structures in the open state, that are often required to see alternate conformations in other proteins^37–41^. The structures generated by AF2 of OSCA1.2 and hTMEM63A/B/C (Fig. 2A) vary on a continuum between open and closed when compared to known open and closed cryo-EM reference structures as measured by their TM score^42^, an indicator of structural similarity between proteins. Interestingly, we note that the highest confidence predictions generated by AF2 are more similar to known open states than closed states (Fig. 2A). The hTMEM63A AF2 predictions at the ends of this continuum resemble the closed structure of hTMEM63A resolved in nanodiscs, and the open structures of OSCA1.2 and TMEM63B respectively (Fig. 2B, S3), with similar results for hTMEM63B and hTMEM63C. The diverse conformers cluster into structurally similar groups (Fig. S4A); with key differences in the TM 3-4 and TM 5-6 helices caused by bending at T523 and T438 (Fig. S4B,D,E), and distinct side chain conformations of W613 which points toward W473 only in closed structures (Fig. S4B,C,F).

**Figure 2.**
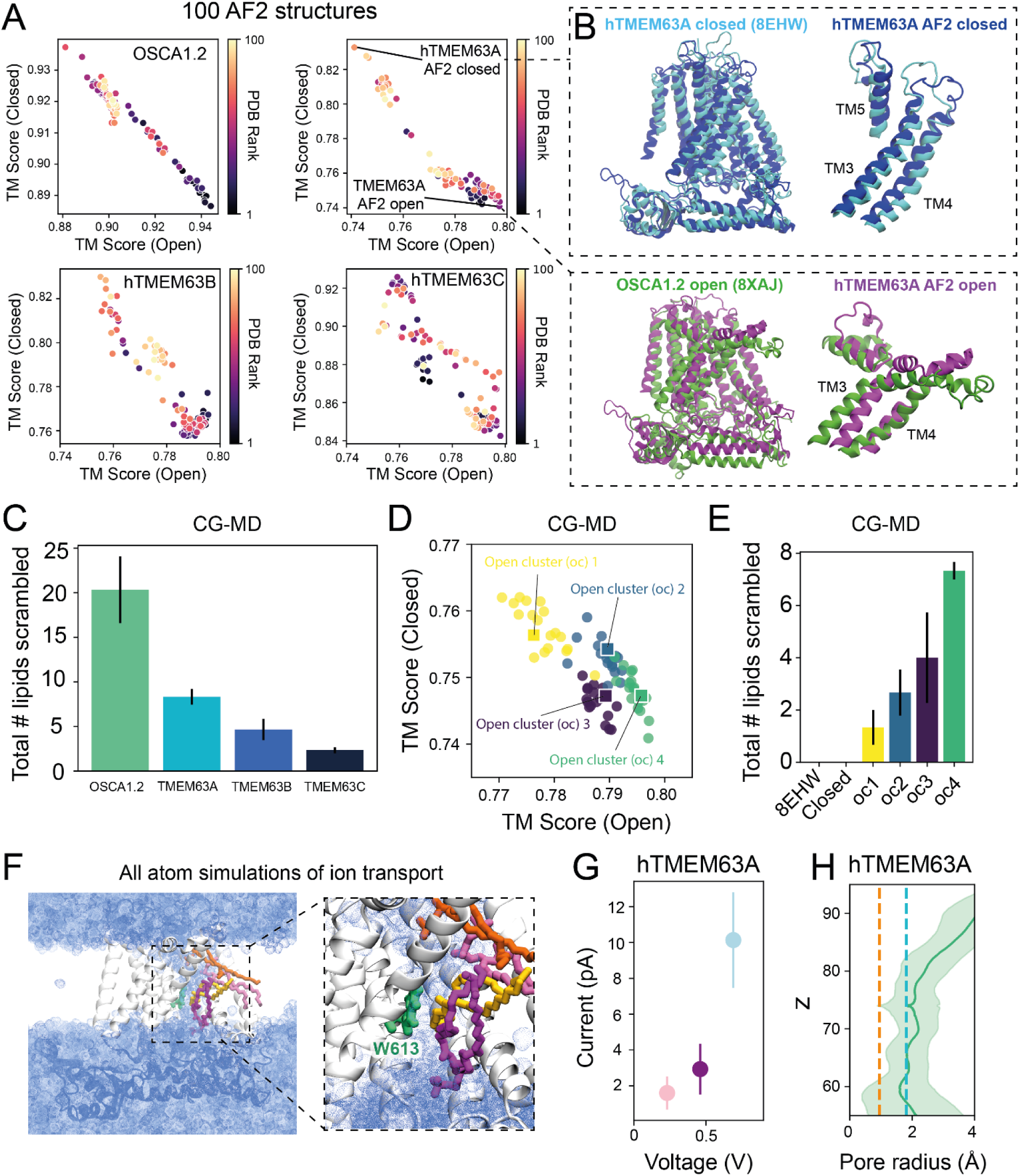
Predicted structures of TMEM63 proteins show intermediate states and an open hydrophilic groove that can scramble lipids and conductions. (A) 100 OSCA1.2, hTMEME63A, hTMEM63B and hTMEM63C structures generated by AlphaFold2 plotted based on similarity (as measured by TM score) to the relevant resolved closed state (OSCA1.2 – PDB ID: 8XNG^5^, TMEM63A – PDB ID: 8EHW^4^, TMEM63B - PDB ID: 8EHX^4^, TMEM63C – PDB ID: 8K0B^18^) vs similarity to the open state of OSCA1.2 (PDB ID: 8XAJ) ^5^. Each structure data point is colored by its AlphaFold2 rank (B) Top: Structural alignment of closed cryo-EM TMEM63A (light blue, PDB ID: 8EHW) and the most closed TMEM63A AF2 prediction (dark blue), zoom in of TM3-5 shown on the right. Bottom: Structural alignment of one cryo-EM OSCA1.2 open monomer (green, PDB ID: 8XAJ) and the most open hTMEM63A AF2 prediction (pink), zoom in of TM3-5 shown on the right. (C) Number of lipids scrambled in 20 μs coarse grained molecular dynamics simulations of AF2 structures of OSCA1.2, hTMEM63A, hTMEM63B, hTMEM63C in the open states. Error bars represent SEM (n=3). (D) 74 structures from the open cluster subdivided into four separate clusters (‘oc1’, ‘oc2’, ‘oc3’ and ‘oc4’) of increasing openness. The central structure of each cluster is indicated by a square data point. (E) Number of lipids scrambled over time in coarse grained molecular dynamics simulations for each TMEM63A open cluster as well as the cryo-EM and AF2 closed states. Error bars are SEM (n=3). (F) Snapshot of atomistic system, with backmapped hTMEM63A shown in grey, pore lipids in orange, light pink, yellow and hot pink licorice. Water is shown as a dotted blue surface. The bottleneck W613 residue is shown in green licorice. (G) A current-voltage plot showing sodium current through the channel at different (230 mV, 460 mV and 690 mV) voltages (shown in pink, purple and blue respectively). Error bars represent SEM (n=3). (H) Pore radius profile for the WT hTMEM63A pore. The solid green line indicates the mean pore radius, with the coloured bands representing one standard deviation of variation (calculated across every frame of triplicate simulations). The dashed orange line indicates the radius of a dehydrated sodium ion. The dashed blue line indicates the radius of a dehydrated chloride ion.

To test whether the predicted hTMEM63 structures facilitate scrambling as seen in the GUV assay (Fig.1F-G) we performed triplicate 20 μs CG-MD simulations of the most open structure of hTMEM63A/B/C as well for the predicted open state of OSCA1.2. Lipids spontaneously translocated through the groove for each these proteins (Fig.2C, Supplementary Video 2) with OSCA1.2 > TMEM63A > TMEM63B > TMEM63C. Lipid translocation happens in both directions in the simulations, but asymmetries in the initial lipid configuration can favor transport in one direction in the early part of the simulation.

Given that hTMEM63A has the highest rate of lipid scrambling in both the experimental assay and the CG-MD simulations, we further clustered the 74 open structures into four groups with increasing levels of “openness” (Fig.2D). While all these structures show a more open hydrophilic groove than the closed state, both the groove width and TM helix tilt increase as we move from open cluster 1 (oc1) to the most open cluster (oc4). Structures from all four can scramble lipids in CGMD simulations (Fig.2E) with a statistically significant difference in flipping rates between the most open (oc4) and least open (oc1) cluster, suggesting that the AF2 generated structures exhibited varying levels of “openness” and may thus represent intermediate conformations.

To investigate whether ions can conduct through the open hTMEM63A groove, we backmapped the end frames of the oc4 CG-MD simulations to atomistic resolution and simulated each for 200 ns with membrane tension applied to keep the channel open. In these simulations a water filled pore remained between the lipids and protein (Fig. 2F) and lipid headgroups were mobile in the groove. Several ion permeation events were observed at membrane potentials of 230, 460 and 690 mV with an estimated unitary conductance of 9.3 ± 2 pS (Fig. 2G), with a preference for Na^+^ permeation over Cl^-^. The current and pore radius is independent of the membrane tension used in the simulation (Fig. S5A,B). This conductance is more than an order of magnitude lower than the experimental or simulated conductance seen for OSCA1.2^5^, in line with the extremely low channel currents recorded from hTMEM63A^2^. The low conductance may be explained by the narrow size of the hTMEM63A pore, with a pore radius of approximately 1-3 Å. A bottle neck is formed by W613 interacting with pore lining lipids (Fig. 2H) requiring ions to partially dehydrate to pass (Fig. S5C). Since the lipid component of the pore is mobile, the size of the pore is more flexible than a pure protein pore and lateral motions of the phospholipids allow the ion to pass through the conduction pathway. Together, this data is consistent with experiments and shows that the AF2 predicted structures can scramble lipids and conduct ions through a lipid lined pore.

### Separate bottleneck residues control ion conduction and lipid scrambling in TMEM63A

Next, we sought to determine what residues control ion conduction and lipid scrambling through TMEM63A. Given the role W613 plays in reducing ion flux in our atomistic simulations, we mutated this residue to asparagine, which eliminates this bottleneck (Fig. 3A). Consequently, the channel conductance is doubled in simulations (Fig.3B). In agreement with this, cell-attached patch clamp recordings also show an approximately two-fold increase in the current density of patches expressing the homologous mutation in mTMEM63A (W613N) compared to WT (Fig. 3C-D), supporting the location of the pore predicted by AF2 and that this residue forms a bottleneck for ion permeation.

**Figure 3.**
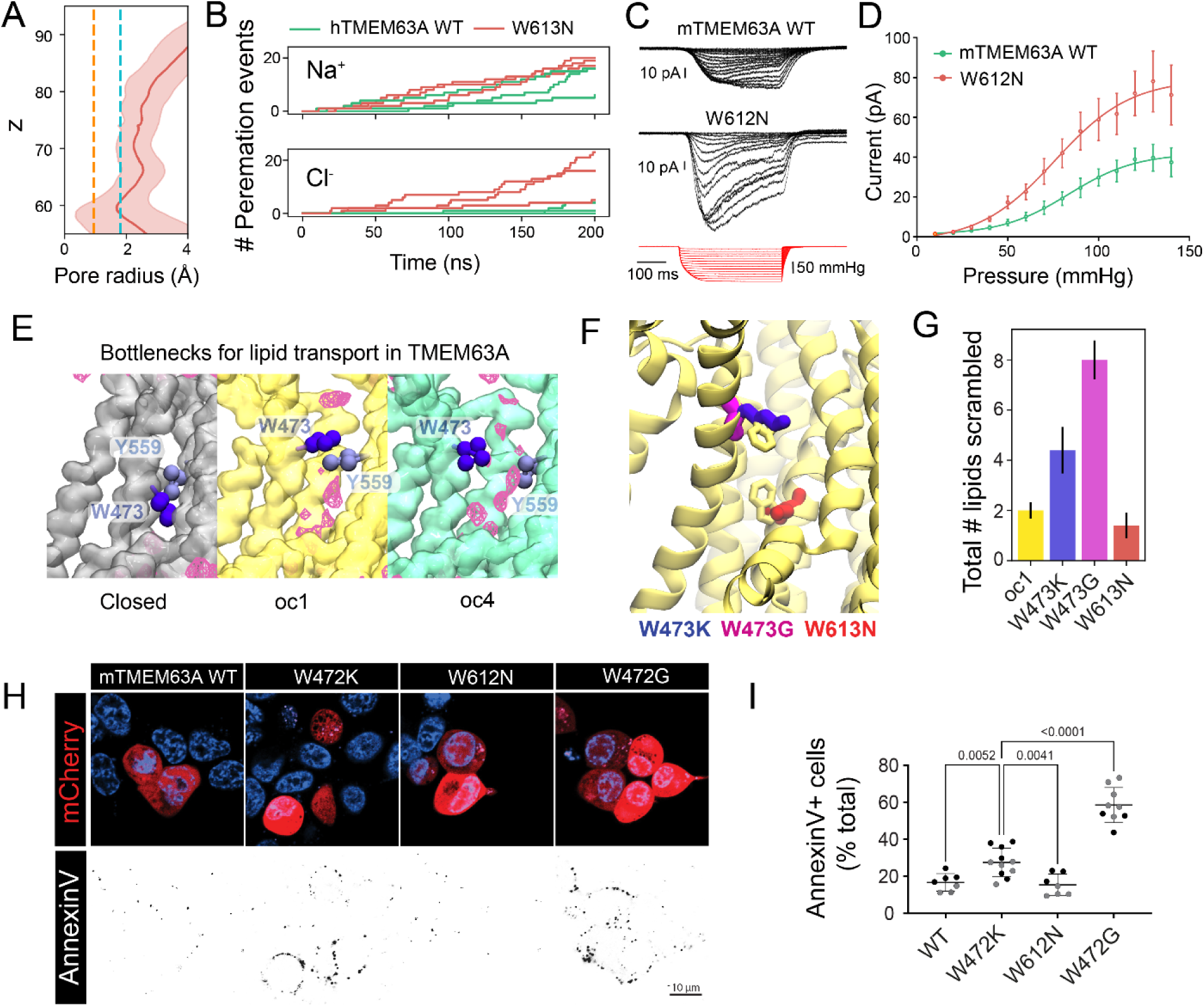
Separate bottleneck residues control ion conduction and lipid scrambling in TMEM63A. (A) Pore radius profile for the W613N hTMEM63A pore. The solid red line indicates the mean pore radius, with the coloured bands representing one standard deviation of variation (calculated across every frame of 3 triplicate simulations). The dashed orange line indicates the radius of a dehydrated sodium ion. The dashed blue line indicates the radius of a dehydrated chloride ion. (B) Conductance vs time graph for sodium (top) and chloride (bottom) ions through the WT (green) and W613N (red) TMEM63A pore. (C) Cell-attached currents at -65 mV holding potential of WT mTMEM63A and W612N mTMEM63A (mouse numbering, W613N in human TMEM63A) overexpressing *Piezo1^-/-^* HEK293T cells in response to negative hydrostatic pressure applied using a high-speed pressure clamp. (D) Peak current plotted versus negative pressure applied [-mmHg] WT mTMEM63A (n=7) and W612N mTMEM63A (n=10). (E) Phosphate density (pink mesh) in hTMEM63A groove in the closed cryo-EM structure, oc1 and oc4 with the side chain beads of bottleneck residues W473 and Y559 shown in VdW. (F) Locations of key residues mutated in this study are shown in liquorice in the TMEM63a oc1 groove. (G) Total number of lipids scrambled in replicate WT and mutant oc1 CGMD simulations of length 40 μs for each system. Error bars represent SEM (n=5). (H) Confocal images of *Piezo1^-/-^*HEK293T cells expressing mTMEM63A WT, W472K (W473 in human TMEM63A), W472G and W612N (W613N in human TMEM63A) labeled with AnnexinV-FITC. (I) Quantification of AnnexinV-FITC positive cells as a percentage of cells transfected with mTMEM63A constructs shown in panel j. Grey and black points reflect clone #1 and #2 of the *Piezo1-/- Tmem16f-/-* double KO cells. Error bars represent SEM (n=7-11) and p-value determined using one-way ANOVA with Tukey’s post hoc test.

Lipids undergoing translocation in CG-MD simulations interact with a number of bulky aromatic residues (W613, W473, Y559 & Y484) that line the groove (Fig. S5D). We find that the residues W473 and Y559 form a gate for lipids near the mouth of the groove which is closed in the cryo-EM and oc1 structures, and prevents or severely limits lipid translocation in all but the most open cluster (Fig.3E, S5E). Mutations W473K and W473G at this gate (Fig. 3F) increase the rate of scrambling in CG-MD simulations with the greatest effect seen for W473G, while mutation of the bottleneck controlling ion permeation, W613N, did not alter lipid scrambling (Fig. 3G, S5H). The mutations have less influence in the most open oc4 structure compared to oc1, as W473-Y559 is not the main bottleneck in this conformation (Fig. S5F,G) which is instead formed by the interaction of Y484 with adjacent residues (e.g. P576) lower down in the groove (Fig. S5F).

To experimentally test the effect of these mutations we turned to a cell based scrambling assay that detects the presence of phosphatidylserine (PS) on the outer leaflet of both *Piezo1^-/-^*and *Piezo1^-/-^ Tmem16f^-/-^* double knockout HEK293T cells expressing the wild-type (WT) mTMEM63A or its analogous mutants. We observed a profound increase in annexinV-FITC labelling as a marker of PS exposure in both cell lines expressing mutations at W472K/G. Consistent with simulations, the W472G mutant increased annexinV labelling the most (Fig. 3H,I). This suggests that these mutants have spontaneous lipid translocation in the absence of an overt exogenous stimulus and are constitutively open. In contrast the W613N mutation does not show significantly different PS exposure compared to WT (15.1 ± 3.8 %; n=6).

### Cholesterol inhibits lipid scrambling in TMEM63A

Results from our experimental scrambling assays show that WT TMEM63A translocates lipids in GUVs but this is not the case in the cellular context. This suggests either; (i) there is a stimulus already present in the GUV assay (perhaps due to the osmotic swelling used to make GUVs) or (ii) factors present in the cell may act to inhibit lipid scrambling. Given that the membrane composition, and in particular the presence of cholesterol, is known to affect currents in TMEM16 and scramblase activity in GPCRs^43,44^, we carried out simulations of the oc4 (most open) structure embedded in two component (POPC/CHOL) bilayers with increasing cholesterol concentrations (5, 10 and 20%) as well as a model ‘complex’ membrane containing POPC, POPS, POPE, PIP_2_ and CHOL. The presence of cholesterol significantly slows lipid scrambling (Fig. 4A). The major bottleneck for lipid permeation in oc4 is at the location of Y484, and this constriction is narrowed in a cholesterol concentration-dependent manner (Fig. 4B). This reduction in scrambling could be due to cholesterol directly binding to the protein or by it altering bulk membrane properties such as membrane rigidity, lipid diffusion or membrane pressure profiles, or by dissipating interleaflet lipid asymmetries present initially as it can freely flip-flop without the aid of a protein. To assess this, we carried out simulations where 20% cholesterol was present but prevented from interacting with the protein (20%-res; Fig. 4A-B). These simulations also exhibited slowed lipid flipping (Fig. 4A) and a narrowed Y484-P576 bottleneck, suggesting that cholesterol slows scrambling indirectly (Fig. 4B).

**Figure 4.**
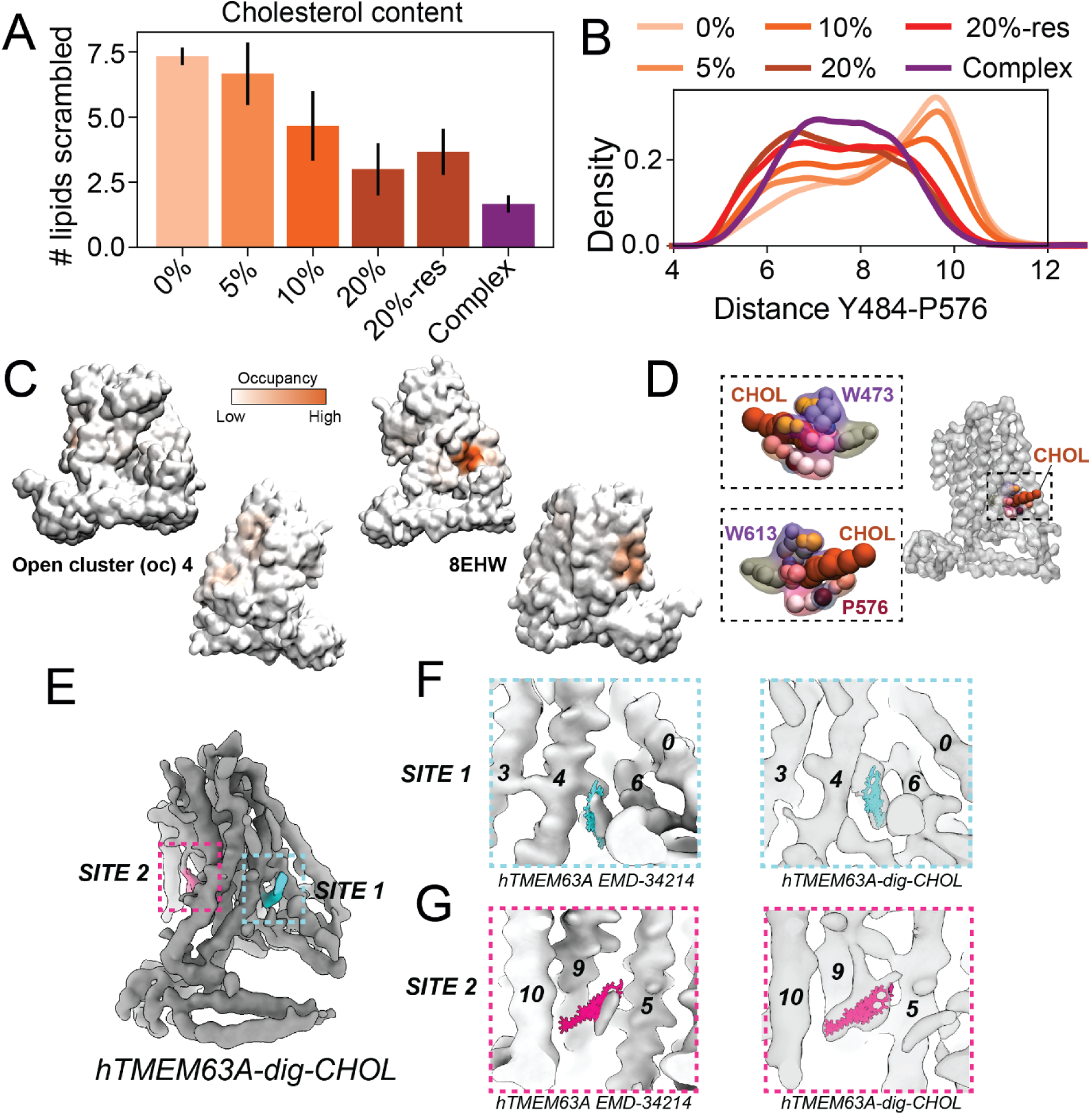
Cholesterol binds hTMEM63A and inhibits lipid scrambling *in silico*. (A) Total number of lipids scrambled through hTMEM63A in 20 μs for membranes containing increasing amounts of cholesterol as well as a system with 20% cholesterol restrained away from the protein (20%-res) and a complex 5 component membrane system containing 20 % cholesterol (Complex). Error bars represent SEM (n=3). (B) Probability distribution of the bottleneck distance between the Y484-P576 sidechains across different membrane systems shown in panel A, indicating a narrowing in the presence of cholesterol. (C) Contact residency surface for cholesterol on the open hTMEM63A structure (AF2 generated, oc4) and the closed hTMEM63A structure (PDB ID: 8EHW), viewed from 2 different angles. (D) Cholesterol binding the closed hTMEM63a groove, shown on the entire protein surface along with close up views from two orientations highlighting the interacting residues. (E) Cryo-EM map (grey) of hTMEM63A with cholesterol (pink and cyan). Two cholesterol binding sites are highlighted in dashed boxes. (F) Zoomed-in views of binding site1 shown in (E). The map is displayed as a transparent surface with a model of cholesterol fitted in (cyan). (G) Zoomed-in views of binding site2 shown in (E). The map is displayed as a transparent surface with a model of cholesterol fitted in (pink).

In addition to slowing scrambling when the channel is open, cholesterol may also inhibit hTMEM63A scrambling through stabilization of the closed state. Analysis of cholesterol interactions in simulations of the closed cryo-EM structure of hTMEM63A (PDB ID: 8EHW) revealed a high residence time binding interaction (Fig. 4C-D) on the groove interface which is not present in the open state. This interaction occurred in two out of three replicate simulations, with cholesterol interacting for the entirety of the 20 μs simulation in one replicate and stable in the final 5 μs of another replicate in an identical pose. At this site, cholesterol interacts mainly with W473, W613 and P576 via the hydroxyl group of cholesterol (Fig. 4D). The presence of this site in the closed state but not in the open state suggests that cholesterol would stabilise the closed state of TMEM63A. To structurally characterize the interaction of cholesterol with hTMEM63A we mixed cholesterol with hTMEM63A in digitonin and carried out cryo-electron microscopy (Table S1, Fig S6). We identified extra density in two specific sites (Fig. 4E). Site 1 is at the pore exit and the end of the groove at the same location seen in the MD simulations, bridging TM4 and TM6B, both of which are critical for groove opening (Fig. 4F). Site 2 is located at the cleft surface (the cleft refers to the central cleft in the OSCA dimer), surrounded by TM5, TM9, and TM10 (Fig. 4G).

To experimentally test the role of cholesterol we turned back to the GUV scrambling assay in which the presence of cholesterol can be controlled. When 20% cholesterol was added to hTMEM63A containing GUVs formed from soy polar lipids, the decline in NBD/Rhod fluorescence ratio was indistinguishable from the empty control in which no protein is present (Fig.5A-C), in contrast to GUVs without cholesterol (Fig.1F-G).

**Figure 5.**
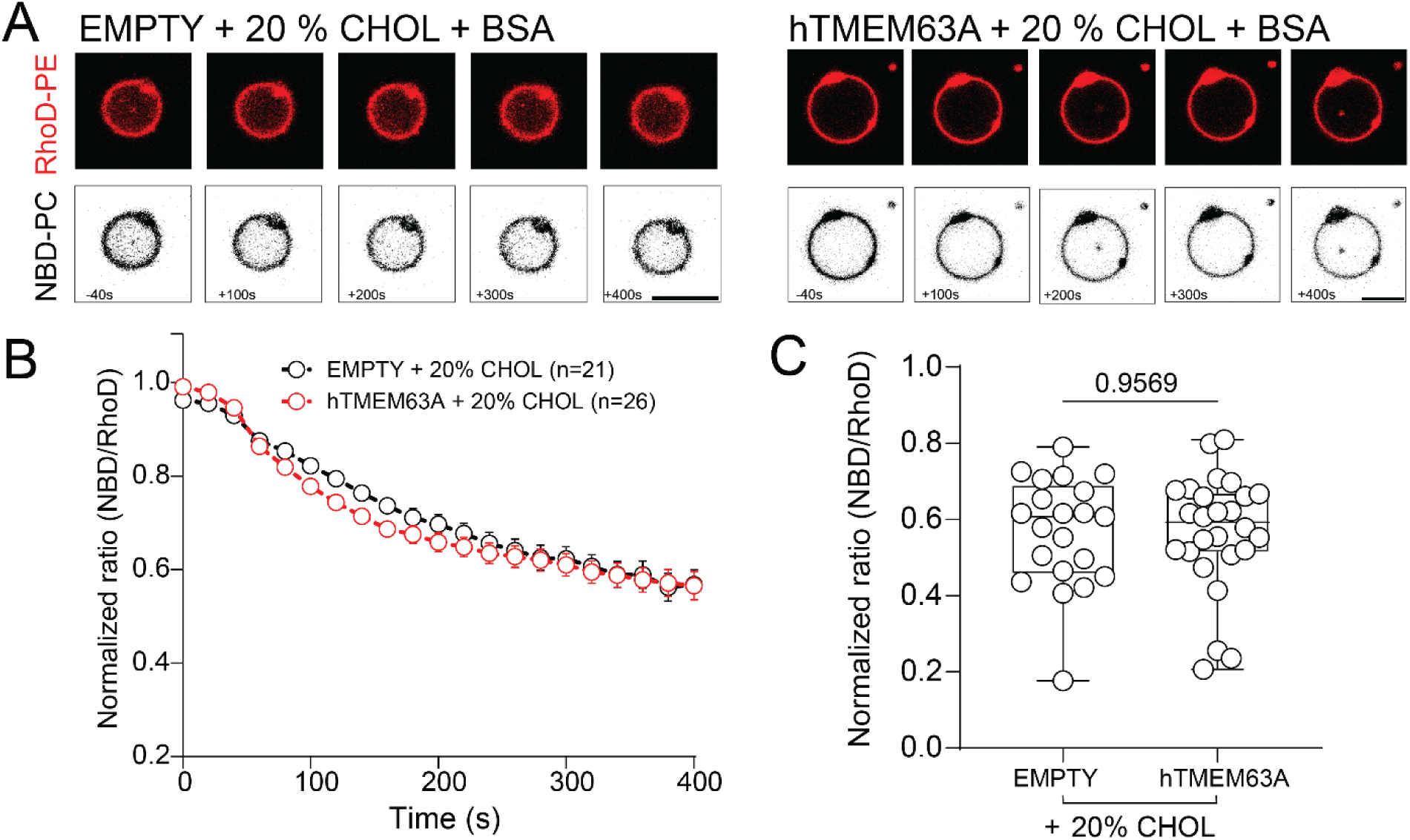
Cholesterol inhibits hTMEM63A facilitated lipid scrambling *in vitro*. (A) Confocal images of soy polar lipid GUVs containing 0.2 % w/w Rhod-PE and 0.8 % w/w NBD-PC and 20 % cholesterol with and without hTMEM63A before and after addition of 4 mg/mL BSA (scale bar = 5 µm). (B) Time course of changes in normalized ratio of NBD/Rhod fluorescence intensity after BSA addition in empty (n=21) and hTMEM63A (n=26) containing GUVs in the presence of 20% cholesterol. Data represents mean ± SEM. (C) Normalized ratio of NBD/Rhod fluorescence intensity 400 s after BSA addition from the data displayed in panel F. Data shown as max to min box and whiskers plots and p-value determined using unpaired Student’s t-test.

### Lipid scrambling in TMEM63A can be mechanically activated

Having shown that OSCA1.1/1.2/2.2 and TMEM63A/B can translocate lipids, and that they are mechanically activated ion channels, we next sought to assess if lipid scrambling could also be mechanically activated. Making use of the fact that hTMEM63A shows no scrambling in cholesterol containing GUVs we developed a method to further increase mechanical force in the cholesterol containing GUVs to unequivocally prove force-induced lipid scrambling by TMEM63 proteins (Fig.6A). We have previously used cyclodextrins (CDs) for functional and structural studies of MS channels^45,46^. Given the influence of cholesterol on hTMEM63A scrambling here we utilised α-cyclodextrin (αCD) which can remove diverse phospholipids from membranes but lacks sufficient internal diameter to accommodate cholesterol^47^. Thus, when applied to our GUVs αCD should increase membrane tension without removing cholesterol.

**Figure 6.**
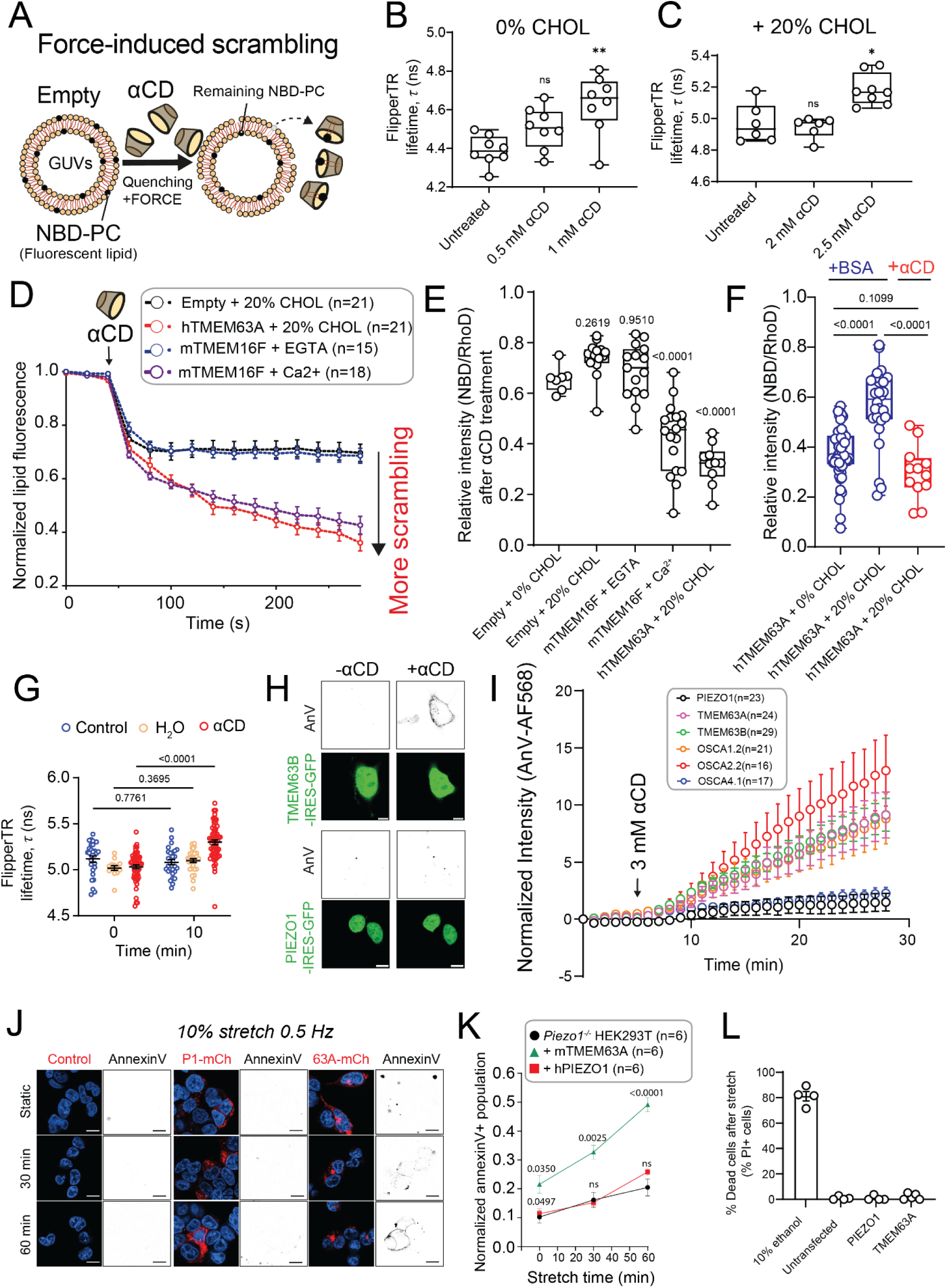
TMEM63A scrambling can be driven by mechanical cues *in vitro* and *in situ*. (A) Model for the novel force-induced lipid scrambling assay using cyclodextrins. (B) Fluorescence lifetime of Flipper-TR incorporated into GUVs made of soy polar lipids with and without αCD. Data shown as max to min box and whiskers plots and p-value determined using one-way ANOVA. (C) Fluorescence lifetime of Flipper-TR incorporated into GUVs made of soy polar lipids supplemented with 20 % cholesterol with and without αCD. Data shown as max to min box and whiskers plots and p-value determined using one-way ANOVA. (D) Time course of changes in normalized ratio of NBD/Rhod fluorescence intensity after αCD addition in empty GUVs containing 20% cholesterol (n=21), and GUVs reconstituted with in the presence of 20% cholesterol (n=21) and mTMEM16F with 2 mM EGTA (n=15), mTMEM16F with 1 mM Ca^2+^ (n=18). Data represents mean ± SEM. (E) Normalized ratio of NBD/Rhod fluorescence intensity 400 s after 2.5 mM αCD addition from the data displayed in panel D. Data shown as max to min box and whiskers plots and p-value determined using one-way ANOVA with Dunnett’s post hoc test. (F) A direct comparison of the normalized ratio of NBD/Rhod fluorescence intensity 400 s after either the addition of BSA or αCD (G) Fluorescence lifetime of Flipper-TR incorporated into HEK293T cells exposed to control conditions, 70% H2O as a hypo-osmotic shock and 5 mM αCD. Data represents mean ± SEM and p-value determined using one-way ANOVA. (H) Representative images of AnnexinV-AlexaFluor55 labelling in HEK293T cells treated with 2.5 mM methyl-β-cyclodextrin before and after the addition of 3 mM αCD. (I) Normalized intensity of annexinV-AlexFluor568 of *Piezo1-/-* HEK293T cells exposed to 2.5 mM methyl-β-cyclodextrin at time point 0 and then 3 mM αCD at the black arrow. (J) Confocal images of control *Piezo1^-/-^*HEK293T cells and the same cells expressing mCherry fused versions of mTMEM63A WT and hPIEZO1 (as a negative control) after 0, 30 and 60 minutes of cyclic uniaxial stretch at 10% strain and 0.5 Hz. AnnexinV-FITC labelling is represented in black (scale bar = 10 µm). (K) Quantification of the normalized amount of transfected cells labelled with annexinV-FITC relative to uniaxial stretch duration. Data points represent mean ± SD (n=6). P value was determined using two-way ANOVA compared to *Piezo1^-/-^* HEK293T cells. (L) % dead cells as indicated by PI staining after 60 min cyclic stretch.

To ensure αCD can increase force within the GUVs we used the reporter of membrane stress Flipper-TR. In simplified systems an increase in tension/stress is reported by Flipper-TR as an increase in the fluorescence lifetime^48^. In GUVs made of soy polar lipids treatment with 1 mM αCD increased lifetime indicative of increased stress within the membrane (Fig.6B). In GUVs made of soy polar lipids supplemented with 20 % cholesterol 2.5 mM αCD was required to generate an increase in fluorescence lifetime of the Flipper-TR probe (Fig.6C).

To see if increased tension could facilitate lipid scrambling, we then applied αCD to increase forces in hTMEM63A GUVs made of soy polar lipids supplemented with 20 % cholesterol. We observed that, much like BSA, treatment with αCD selectively quenched NBD-PC over Rhod-PE, meaning we could mechanically induce scrambling with αCD alone. In contrast to data with BSA where cholesterol prevented hTMEM63A scrambling (Fig.5) we observed that under force in the presence of αCD the relative intensity of NBD/Rhod decreased markedly in GUVs containing hTMEM63A, much more than seen in empty liposomes (Fig.6D,E). CDs asymmetrically remove lipids from the GUVs so to rule out this process alone was driving scrambling we treated mTMEM16F containing GUVs with αCD in the presence of EGTA. We then observed no evidence of lipid scrambling (Fig.6D,E). Under these same conditions, we observed scrambling from mTMEM16F when Ca^2+^ was present. This very clearly and robustly showed that αCD did not drive scrambling unless a lipid translocation pathway was present. When we compared cholesterol containing GUVs treated with αCD against BSA, it was clear that TMEM63A only exhibited scrambling if increased force was applied (Fig.6F).

As purified and reconstituted hTMEM63A protein displayed mechanically induced scrambling and given conflicting previous results on whether the WT TMEM63 proteins may represent scramblases we sought evidence for TMEM63A-induced scrambling *in situ* ^34,49^. Previous work suggested that the removal of cholesterol alone by methyl-β-cyclodextrin (mβCD) can drive TMEM63B scrambling^33^. This ignores the essential fact that CDs of all types can increase membrane tension ^45,50^. To prove this conclusively, we again turned to αCD, which cannot remove cholesterol. Using the Flipper-TR probe we showed treatment with αCD increases the lifetime of the probe well above that of a significant hypo-osmotic shock (70% H_2_O) in HEK293T cells (Fig.6G). This is consistent with increased membrane stress. We also tried the same for mβCD but noted that the Flipper-TR probe itself was removed from the membrane by mβCD.

Next, we wanted to show that low doses of mβCD (2.5 mM) can’t drive scrambling but that the subsequent addition of αCD (which can’t remove cholesterol but can increase membrane forces) drives significant scrambling in cells expressing OSCA/TMEM63 proteins (Fig.6H-I). As a negative control we also used OSCA4.1, a protein we found cannot escape intracellular compartments in HEK293T cells (Fig.S7). In this scenario we saw little scrambling after mβCD (2.5 mM) addition but after subsequent αCD addition we saw robust annexinV labeling in cells expressing OSC/TMEM63 proteins (Fig.6H-I). This data is consistent with mβCD working through changes in membrane stress rather than only through cholesterol.

To provide further context for mechanically induced scrambling we expressed mTMEM63A *in Piezo1^-/-^* HEK293T cells then exposed these cells to uniaxial cyclic stretch with 10 % strain at 0.5 Hz for 0, 30 and 60 minutes. Here we also expressed hPIEZO1 as a negative control. We saw a profound increase in PS exposure indicated by annexinV-FITC labeled cells in the mTMEM63A expressing group that was not present in untransfected *Piezo1^-/-^* HEK293T cells and much higher than the group of cells expressing hPIEZO1 (Fig.6J-K). PIEZO1 is much more sensitive to mechanical force and has a significantly higher unitary conductance than TMEM63A so this data is congruent with PS scrambling being driven by mechanically activated TMEM63A. We saw similar effects when OSCA1.1 and 1.2 were transiently expressed in the same cell type (Fig.S8). Cells stretched for this period did not label with propidium iodide (PI) (that enters and binds to DNA in dead cells, causing them to fluoresce) showing that they were not dying, providing support for the specificity of annexin-V labelling (Fig.6L).

### TMEM63 influences membrane remodeling and resilience

An obvious question is; what could the relevance of the mechanically activated scramblase activity of TMEM63A/B be in a cellular environment? While over-expressed TMEM63A in HEK293T cells does reach the plasma membrane, making it possible to record mechanically induced currents, native TMEM63A has been shown to be localized to lysosomes in drosophila and mice^51^. In this organelle TMEM63A is involved in membrane morphological changes. So, we next asked the question of whether hTMEM63A influences membrane morphology under mechanical force (Fig.7A-C). When empty GUVs made of soy polar lipids were exposed to high doses of αCD, ∼60% broke within 100 seconds with similar patterns observed with GUVs containing hTREK2 or the bacterial mechanosensitive channel MscS (Fig.7B). However, in the presence of the scramblases nhTMEM16 or hTMEM63A the liposomes tended to shrink in size (Fig.7A,C) with >80 % of GUVs surviving more than 300 s. In GUVs that contained hTMEM63A, force induced scrambling overwhelmingly induced membrane tubule formation that was rarely identified in any other experimental condition (Fig.7A,D). This suggests mechanically induced scrambling through hTMEM63A can drive morphological changes in membranes in response to high mechanical forces and protect them from rupture.

**Figure 7.**
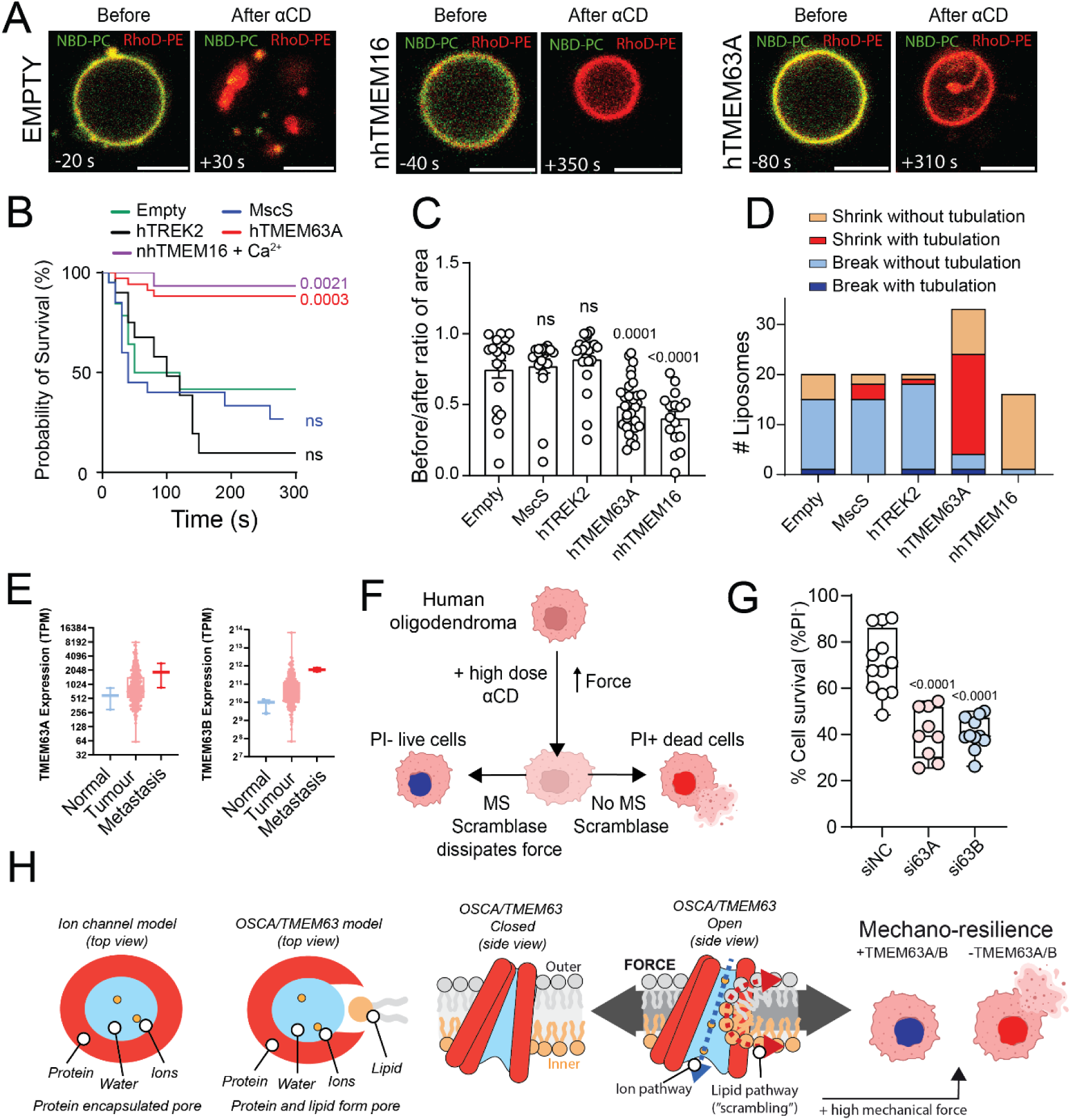
hTMEM63A influences membrane morphology and mechano-resilience. (A) Confocal images of soy polar lipid GUVs containing 0.2 % w/w Rhod-PE and 0.8 % w/w NBD-PC and 20 % cholesterol with and without hTMEM63A before and after addition of αCD (scale bar = 5 µm). (B) Kaplan-Meier curve of GUV survival under force after the application of αCD in empty GUVs (n=20) and GUVs containing hTREK2 (n=20), MscS (n=20), nhTMEM16 (n=16) and hTMEM63A (n=33). P-value determined using log rank Mantel-Cox test. (C) Quantification of the 2D area of the liposomes before and after the addition of αCD in all the GUV conditions shown in panel B. (D) Categorization of the response of GUVs in panel B after the addition of αCD. (E) Expression level of TMEM63A and TMEM63B in normal, tumour and metastic cervical cancer. (F) Cartoon depiction of the in vitro mechano-resilience assay. (G) Cell survival of HOG cells treated with siRNA negative control (siNC), siRNA for TMEM63A and TMEM63B subsequently exposed to X mM αCD. Data shown as max to min box and whiskers plots and p-value determined using one-way ANOVA. (H) Model for open state of OSCA1.2 and subsequent mode for lipid scrambling through TMEM63B and its influence on mechano-resilience.

### TMEM63A contributes to cellular mechano-resilience in native cells

Given that previous work has suggested that TMEM63B resides at the plasma membrane and that TMEM63A can reach the plasma membrane in some cell types^52^ we wondered if lipid scrambling by these proteins has additional physiological functions beyond organelle morphology. We noticed that TMEM63A was highly expressed in some metastatic cancers over “normal” cells while TMEM63B was highly expressed in almost all metastatic cancers. Metastatic cancer cells are required to resist high mechanical forces selecting for those with increased mechano-resilience (Fig 7E, Fig S9). Thus, to test if TMEM63 proteins could affect cellular mechano-resilience we turned to a cancer cell line that expresses both TMEM63A and TMEM63B, namely human oligodendroglioma (HOG), noting that TMEM63A can reach the plasma membrane in native oligodendrocytes^52^. We treated HOG cells with non-targeting siRNA, TMEM63A or TMEM63B siRNA then exposed them to a high dose of αCD to induce rupture in ∼20 % of the cell population (Fig.7F-G). We observed that knocking down either TMEM63A or TMEM63B increased the number of cells rupturing as reported by propidium iodide staining, consistent with a protective effect of both proteins at native protein levels in a relevant cell line.

## Discussion

In a recent study, cryo-EM structures and MD simulations showed that the ion conduction pathway in the mechanically gated OSCA1.2 ion channel occurs through a lipid-lined pore ^5^. Subsequent papers have suggested that members of both the OSCA and TMEM63 protein families have scramblase capability.^33–35,49^ However, in all these reports it is unclear whether the WT OSCA/TMEM63 proteins have scramblase capability ^34^ or what the true native stimulus for the scramblase activity is *in situ* ^49^. We probed the scramblase activity of TMEM63/OSCA channels using a combination of *in vitro* and *in situ* assays and computational techniques providing compelling evidence that WT members of both the mammalian TMEM63 and plant OSCA ion channel families possess the ability to scramble lipids consistent with the open groove model for lipid scrambling.

Using AlphaFold2 we predicted open structures of proteins within the OSCA/TMEM63 family and simulated these structures alongside the experimentally resolved activated state of OSCA1.2. CG simulations revealed that TMEM63A/B/C and OSCA1.2 all facilitate lipid translocation in the open state. By simulating AF2 generated intermediate states of TMEM63A, we identified several aromatic residues (W473, Y559, W613 and Y484) which form bottlenecks within the open groove. We identified bottleneck residues for ion permeation (W613) and lipid translocation (W473). Consistent with this, simulations and experiments show mutations at the W473 position increased rates of lipid scrambling and hydrophilic substitutions at W613 boost ion conductance. The very narrow, lipid lined pore with an average radius of 2 Å may explain the extremely small unitary conductance of TMEM63A observed in experiments^2,4^, in contrast to large currents which can freely pass through the larger pore (∼ 4 Å) in the related OSCA channels^5^. This study highlights AlphaFold2’s ability to generate diverse conformations of an ion channel without the need for advanced sampling techniques^37–41^. The importance of potential coupling of ion currents to scrambling remains to be determined, especially if both are physiologically relevant or if one is the byproduct of the other. For example, it is not yet clear if the small ion currents with slow kinetics seen for TMEM63A and OSCA2.2 are physiologically relevant or if this is a ‘leak’ current required to enable lipid translocation.

We showed that cholesterol slowed lipid scrambling via the open structure, likely through modulation of bulk membrane properties. Additionally, it binds tightly to the groove in the closed state, likely stabilizing it and increasing the threshold required for mechanical activation. Interestingly, this binding location is the same as was identified for a phospholipid, assigned as phosphatidylcholine in a structure of TMEM63A (PDB ID: 8GRS) solved in detergent^19^ . Our new Cryo-EM structures identified two sites with density consistent with cholesterol in hTMEM63A including this previously identified lipid binding site. In the homologous TMEM16 family, this is also the location of binding for calcium, which is required for channel activation^23,53,54^. Given that cholesterol itself can freely flipflop between leaflets without the aid of a protein, it can dissipate the pressure gradient and hence remove the driving force for lipid flipping. Reduction of activity by cholesterol could be a common inhibition mechanism across mammalian scramblases. For example, GPCRs can scramble lipids when reconstituted in proteoliposomes but not within the context of cellular membranes containing cholesterol^55,56^. MD simulations showed that cholesterol slows phospholipid translocation through the GPCR groove by occluding it and being flipped itself^43^. Additionally, while the effect of cholesterol on TMEM16 scramblases has yet to be investigated, ion currents are inhibited by cholesterol in TMEM16A, a chloride channel in the same family^57^ while TMEM16F is able to scramble lipids in liposomes containing 20% cholesterol^53^. While the mechanisms of cholesterol-mediated inhibition on scramblases awaits further investigation, this suggests that cholesterol and other lipids (such as the phosphoinositides that modulate TMEM16F)^58^ may limit scramblase activity within specific cellular environments. More broadly, our findings support the idea that the activity of scramblases and mechanically gated ion channels are both critically regulated by membrane components^58–62^.

Our data provides compelling evidence that OSCA/TMEM63 proteins serve a dual function as mechanically activated ion channels and lipid scramblases. Lipid scrambling is observed only during simulations of OSCA/TMEM63 channels in the open states similar to atomistic simulations that have been employed to provide structural insights into lipid scrambling via TMEM16 proteins^28–30^. However, it is important to note that much of the experimental and structural data in TMEM16 proteins suggests an alternate mechanism such as out of groove scrambling^63^. We show that in contrast to typical ion channel proteins in which the ion conducting pathway is formed exclusively by the protein, in the family of TMEM63/OSCA proteins the pore wall is partly comprised by lipid molecules (Fig. 7H). Under the influence of mechanical forces, a hydrophilic groove opens on the side of these proteins, allowing a belt of lipids to form between the membrane leaflets. The lipids themselves do not fully enter the groove, due to the balance of hydrophobic and hydrophilic interactions with the protein and create a water and ion conducting pore. Thus, intuitively we suggest that the size of the groove, the depth to which the lipids penetrate and the ability of lipids to translocate between the bilayers differs among the family to yield a diversity of ion currents and lipid scrambling rates.

Stimulating GUVs with αCD increased tension and stimulated hTMEM63A induced lipid scrambling. This was supported by uniaxial stretch experiments showing TMEM63A caused PS exposure *in situ*. Together this data strongly supports the idea that TMEM63A acts as a mechanically activated scramblase, raising the question of what the physiological role of such proteins could be. We showed in GUVs under high force TMEM63A caused profound tubulation and membrane remodeling not seen in control conditions. It is possible TMEM63A/B could play a role in plasma membrane remodeling or in organelle biogenesis during which membranes undergo significant mechanical stress with directional transport dissipating asymmetries in monolayer stress. This is supported by the fact that the loss of TMEM63A influences lysosome morphology^51^. GUVs containing TMEM63A also survived significantly better under higher mechanical forces. We showed that this is also true in the cellular environment where native levels of TMEM63A/B provide protection against high mechanical force. We thus suggest that TMEM63A/B and their mechanically-induced scrambling could play a role in membrane mechano-resilience. Along these lines both TMEM63A and TMEM63B are more highly expressed in metastatic cancer cells than normal cells. Metastatic cells are required to be mechano-resilient^64^ and thus the mechanisms underlying this are potential anti-cancer targets. In addition, recent studies have shown that phospholipid asymmetries between bilayer leaflets are enabled by the presence of cholesterol that can flip-flop between leaflets^65^ suggesting that alteration in cholesterol levels (naturally or by addition of mβCD) can affect the integrity of the membrane. However, the presence of scramblases can counter this, allowing phospholipids to move to the outer leaflet to equilibrate the membrane and prevent rupture, consistent with our results (Fig 7G) and the fact that scrambling can rescue rat basophilic leukemia cells under mechanical stress in the absence of Ca^2+^ induced scrambling^65^. A mechanically-activated scramblase would enable spatio-temporal control of scrambling to resist high mechanical forces. The activation would then subside as the force dissipates due to the scrambling, making these proteins perfectly suited to this role. This also gives a functional explanation for why cholesterol inhibits lipid scrambling: allowing phospholipid asymmetries to develop in high cholesterol conditions while maintaining resilience to cholesterol depletion. This provides further support that mechanically induced scrambling from TMEM63/OSCA proteins may be involved in mechano-resilience, allowing cells to survive under damaging mechanical forces (Fig 7H right).

In conclusion, our computational, structural, and functional data suggests that OSCA/TMEM63 channels are the first identified lipid scramblases activated by mechanical stimuli. This mechanically induced scramblase is likely linked to the mechano-resilience of biological membranes, allowing cells to survive under otherwise noxious mechanical stimuli. Future research will undoubtedly unveil details of the physiological and or pathological roles of these newly identified mechanically activated scramblases.

## Methods

### AlphaFold2 structure generation and analysis

To obtain diverse structural conformers of TMEM63A in the absence of an experimentally resolved open structure, the hTMEM63A (UNIPROT ID: O94886), hTMEM63B (UNIPROT ID: Q5T3F8), hTMEM63C (UNIPROT ID : Q9P1W3) and atOSCA1.2 (UNIPROT ID: Q5XEZ5) sequences were used to generate multiple sequence alignments for structure prediction by Colabfold.^37,66^ 100 structures were predicted for each protein without relaxation or modification to the multiple sequence alignment. TM scores were calculated using TMalign with loop residues omitted.^42^ Cluster analysis of the 100 structures was performed using gmx cluster. Structural analyses (RMSF, inter-residue distances, side chain conformations) were carried out using python scripts using the MDAnalysis package.^67^ VMD was used to visualize structures and prepare figures.^68^

### Coarse grained (CG) molecular dynamics simulations

All coarse-grained simulations were carried out using GROMACS 2023^69^ with the Martini 2.2 forcefield^70^ and prepared using bash and python scripts adapted from the MemPROTMD pipeline^71^ which utilizes martinize for resolution transformation and insane for membrane embedding, solvation and addition of 0.15 M NaCl.^70^ Each system was energy minimized and equilibrated in the NPT ensemble for 1 ns with a timestep of 10 fs and 0.75 ns for a timestep of 15 fs. During equilibration, the Berendsen barostat was used to maintain a pressure of 1 bar with semi-isotropic conditions.^72^ An elastic network and position restraints of 1000 kJ mol^-1^ nm^-2^ were applied to the protein backbone beads in all simulations. Electrostatics were treated using the reaction-field method with a dielectric constant of 15, and a Coulomb cutoff of 1.1 nm. A cutoff of 1.1 nm was applied for van der Waals interactions.

During production simulations, a pressure of 1 bar was maintained using a Parrinello-Rahman barostat with semi-isotropic conditions. The temperature was maintained at 310 K using a v-rescale thermostat. A timestep of 20 fs was used in all production simulations.

To assess scrambling activity, CG simulations of closed TMEM63A (PDB ID: 8EHW^4^), closed TMEM63B (PDB ID: 8EHX), closed TMEM63C (PDB ID: 8k0b), closed OSCA1.2 (PDB ID: 8XNG), open OSCA1.2 (PDB ID: 8XAJ^5^), open nhTMEM16 (PDB ID: 4WIS^54^), the most closed and open AF2 TMEM63A, TMEM63B, TMEM63C and OSCA1.2 predictions, and the central structure belonging to each TMEM63A open subcluster (‘oc1’, ‘oc2’, ‘oc3’, ‘oc4’) were set up. Each structure was prepared according to the protocol above and embedded into POPC membranes. Production simulations were carried out in triplicate for either 20 or 40 us.

To investigate the effect of mutations (W473G, W473K, W16N) on the rate of lipid scrambling, mutations to the atomistic oc4 and oc1 structures were introduced using the biobb structure checking tool.^73^ The resulting structures were energy minimized to remove any steric clashes before being embedded into a POPC membrane. Production simulations were carried out 40 μs with 5 replicates for each mutant.

To assess the effect of membrane composition on lipid scrambling rates, the oc4 structure was additionally simulated in 5, 10, 20% cholesterol as well as a model 5-component membrane with 10% POPS, 5% POP2, 30% POPC, 35% POPE and 20% CHOL in the inner leaflet and 65% POPC, 15% POPE and 20% CHOL on the outer leaflet. These simulations were set up and carried out following the above protocol, for 20 μs in triplicate.

### All atom molecular dynamics simulations

All atomistic simulations were carried out using GROMACS 2023^69^ with the CHARMM36 forcefield.^74^

All atom molecular dynamics simulations were initiated by backmapping the final frame of four oc4 CG simulations using CG2AT.^75^ The backmapped systems were subjected to a short energy minimization, followed by a 5-step equilibration protocol spanning 11.25 ns in total in which restraints on the protein backbone (2000 kJ mol^-1^ nm^-2^, 1000 kJ mol^-1^ nm^-2^, 500 kJ mol^-1^ nm^-2^, 500 kJ mol^-1^ nm^-2^, 10 kJ mol^-1^ nm^-2^), sidechain (2000 kJ mol^-1^ nm^-2^, 1000 kJ mol^-1^ nm^-2^, 500 kJ mol^-1^ nm^-2^, 0 kJ mol^-1^ nm^-2^, 0 kJ mol^-1^ nm^-2^) and lipid headgroups (1000 kJ mol^-1^ nm^-2^, 500 kJ mol^-1^ nm^-2^, 500 kJ mol^-1^ nm^-2^, 500 kJ mol^-1^ nm^-2^, 0 kJ mol^-1^ nm^-2^) were gradually reduced. Periodic boundary conditions were applied in all directions.

Surface tensions of 10 to 60 mN/m were applied in the xy-direction using the Berendsen barostat to keep the channel open during production simulations. Using isotropic pressure coupling, the pressure in the z-direction was maintained at 1 bar..^72^ The temperature was maintained at 310 K using the Nose-Hoover thermostat.^76^ Production simulations used a timestep of 2 fs. Hydrogen bonds were constrained using the LINCS algorithm.^77^ Electrostatic interactions were treated using the Particle Mesh Ewald algorithm with a cut off of 12 Å.^78^A van der Waals radius of 12 Å was used.

### Coarse-grain Simulation Analysis

Analysis of coarse grain simulations are performed across all frames of each simulation trajectory strided such that each frame represents 1 ns of simulation time. Lipid scrambling events in coarse-grained simulations were counted using the scramblyzer package.^79^ Volume density maps, inter-residue distances and occupancy values were calculated using python scripts using the MDAnalysis package.^67^ Probability density plots showing cholesterol or POPC in the groove were calculated by extracting the z positions of PO4 or ROH beads respectively found within 7 Å of groove lining residues in each frame. Distribution plots showing inter-residue distances between groove bottleneck residues used the outermost sidechain bead for each residue (SC4 for TRP, SC3 for TYR, SC2 for PRO, LYS, GLU). Cholesterol binding occupancies were calculated per residue by computing the proportion of simulation time for which any bead belonging to cholesterol could be found within 7 Å of the residue. Cholesterol binding residencies were used to identify stable, long term binding events and calculated the maximum proportion of simulation time for which one specific cholesterol molecule formed an interaction with the residue.

### All-atom Simulation Analysis

Analysis of atomistic simulations are performed across all frames of each simulation trajectory strided such that each frame represents 0.1 ns of simulation time. Ion conduction was analysed using a python script which considers a permeation event to be when an ion moves from a z coordinate of < 70 Å (bottom of the pore) to a z coordinate of > 90 Å (top of the pore), or in the other direction, while remaining within 7 Å of groove lining residues.Pore radius profiles were calculated using hole2,^80^ using the MDAnalysis wrapper,^67^ with both protein and POPC lipids considered in the calculation. Ion coordination numbers were calculated by identifying the number of non-hydrogen atoms belonging to water, lipid or protein found within 3.1 Å of a sodium atom for any frame in which the ion was undergoing permeation.

### Cell culture and transfection

*Piezo1^−/−^* HEK293T cells were a gift from Dr. Ardem Patapoutian (The Scripps Research Institute, La Jolla, CA, United States). HEK293T cells were transfected with polyethylenimine (PEI) as previously described.

### Electrophysiology

Cells were plated on 35 mm dishes for patch clamp analysis. For cell-attached patch clamping configuration, the extracellular solution was: 90 mM K+Aspartate, 50 mM KCl, 2 mM MgCl_2_ and 10 mM HEPES (pH 7.2) adjusted with KOH in order to zero the membrane potential. The pipette solution contained 140 mM NaCl, 2 mM CaCl_2_, 2 mM MgCl_2_, 3 mM KCl and 10 mM HEPES adjusted to pH 7.2 using NaOH. Negative pressure was applied to patch pipettes using a High-Speed Pressure Clamp-1 (ALA Scientific Instruments, Farmingdale, NY, United States) and recorded in millimetres of mercury (mmHg) using a piezoelectric pressure transducer (WPI, Sarasota, FL, United States). Borosilicate glass pipettes (Sigma-Aldrich) were pulled with a vertical pipette puller (PP-83, Narashige, Tokyo, Japan) to produce electrodes with a resistance of 2.0-3.1 MΩ. The currents for mTMEM63A were amplified using an AxoPatch 200B amplifier (Molecular Devices, LLC), and Data were sampled at a rate of 10 kHz with 1 kHz filtration and analyzed using pCLAMP10 software (Molecular Devices, 509 LLC).

### Protein expression and purification

The human TMEM63A gene, was synthesized by GeneWise and cloned (Vazyme ClonExpress II One Step Cloning Kit) into the pEG BacMam vector with a C-terminal GFP tag and a 3C cleavage site between them. The TMEM63A pEG BacMam plasmids were transformed into DH10Bac cells (Weidibio) to generate bacmids. These bacmids were transfected into Spodoptera frugiperda 9 (Sf9) cells using Fugene HD (Promega) in ESF921 medium (Expression Systems) to produce recombinant baculovirus. HEK293F cells deficient in N-acetylglucosaminyltransferase I (GnTI-) were cultured in SMM293-TII Expression Medium (Sino Biological) in a Herocell C1 orbital shaker incubator (Radobio) at 37 °C, 110 rpm, and 8% CO2. When cell density reached 2.5 × 106 cells/ml, P2 baculovirus was added at a 1:20 volume ratio, and the cells were incubated at 37 °C for an additional 20 hours. Following this, 10 mM sodium butyrate was added to enhance protein expression at 30 °C for another 48 hours. Cells were harvested by centrifugation at 3,200 × g for 20 minutes using a JLA 8.1000 rotor (Beckman Coulter) and immediately processed for protein purification.

Cells were resuspended and lysed in lysis buffer containing 30 mM Tris-HCl (pH 7.9), 300 mM NaCl, 2.5 mM MgCl₂, 0.5 mM CaCl₂, 1 mM DTT, 2 μg/mL DNase I, 1 μg/mL aprotinin, 1 μg/mL leupeptin, 1 μg/mL pepstatin, 20 μg/mL trypsin, 1 mM benzamidine, 1 mM PMSF, along with 1% (w/v) DDM and 0.1% (w/v) CHS (Anatrace). The cells were incubated at 4 °C for 2 hours with stirring to solubilize the membranes. The solubilized material was then centrifuged at 30,000 × g for 30 minutes to remove cell debris. 2ml of homemade GFP-nanobody-coated NHS-activated resin (Smart-lifesciences) were added to the supernatant and incubated for 2 hours at 4 °C. The beads were collected by centrifugation and transferred to a gravity flow column. They were washed with 50 mL of wash buffer containing 30 mM Tris-HCl (pH 7.9), 150 mM NaCl, 0.1% (w/v) digitonin, and 1 mM DTT, then incubated with 3C protease for 3 hours at 4 °C to remove the GFP tag. The flow-through and two column volumes of wash buffer were collected and concentrated using Amicon Ultra 15 mL 30 kDa cut-off centrifugal filters (Millipore Sigma). The samples were further purified by size-exclusion chromatography on a Superose 6 10/300 Increase column (Cytiva) connected to an AKTA PURE 25 (Cytiva) or SDL-30 (Sepure Instruments) equilibrated with 30 mM Tris-HCl (pH 7.9), 150 mM NaCl, 0.05% (w/v) digitonin, and 1 mM DTT. Peak fractions were concentrated to a final concentration of 6 mg/mL using a 30 kDa cut-off and immediately prepared for cryo-EM sample freezing. The protein concentration was measured with a spectrophotometer NP80 (Implen). Cholesterol (Sinopharm) was solubilized in ethanol at a concentration of 50 mM, and a final concentration of 1 mM cholesterol was added to the TMEM63A-digitonin mixture for cryo-EM sample freezing.

### Cryo-EM sample preparation and data collection

200 mesh 2/1 Au grids (Quantifoil) were glow discharged for 30 seconds. 4 uL of the TMEM63A-digitonin-cholesterol solution (6 mg/mL TMEM63A with 1 mM cholesterol) was applied to the grids and rapidly vitrified in liquid ethane using a Vitrobot Mark IV (Thermo-Fisher). The vitrification parameters included a waiting time of 10 seconds, a blot time of 1 second, and a blot force of -2, all maintained at 4 °C with 100% humidity.

Data were collected using a 300 kV Titan Krios G4 microscope (Thermo-Fisher) equipped with a Biocontinuum K3 Direct Electron Detector. Micrographs were obtained at a magnification of 81,000x, corresponding to a physical pixel size of 0.83 Å per pixel. The electron dose rate was set to 20.0 electrons per pixel per second, with a total exposure time of 2.2 seconds and a total dose of 63.8 electrons per Å² distributed across 40 frames. A Gatan GIF imaging filter was employed with a slit width of 20 eV. Automated data collection was performed in EPU software ^81^ using the image-shift method, with a defocus range of -1.4 to -2.4 μm.

### Cryo-EM data processing and model building

A total of 15,600 movies from three datasets were imported into RELION-3.1 ^82^, motion-corrected, and electron-dose-weighted using MotionCorr2 ^83^. CTF parameters were estimated with CTFFIND4 ^84^.

Particles were picked using Gautomatch (https://www2.mrc-lmb.cam.ac.uk/download/gautomatch-053/) with templates and extracted from micrographs into 260 x 260-pixel boxes, downsampled to 128 x 128-pixel boxes. These particles were submitted to three rounds of 2D classification in RELION. Particles in 2D classes with sufficient quality were selected, re-extracted into a 420 x 420-pixel boxes, downsampled to 210 x 210-pixel boxes, and submitted to multiple rounds of CryoSPARC ^69,85^ ab initio reconstruction. Particles selected from clear 3D classes were re-extracted into a 420 x 420-pixel boxes at 0.83 Å per pixel. Polished particles in RELION were then submitted to multiple rounds of CryoSPARC heterogeneous refinement using tow bad references and four good references generated from ab initio reconstruction. A total of 202,829 particles from the best classes were subjected to non-uniform refinement in CryoSPARC, resulting in a map at 4.4 Å resolution.

PDB-8GRS was used as the initial model, fitted as a rigid body into the cryo-EM density map using UCSF Chimera ^86^ and then manually adjusted in Coot ^87^. The adjusted model was subjected to iterative rounds of real-space refinement in PHENIX ^88^.

### Reconstituting proteins into liposomes

Purified proteins were reconstituted into giant unilamellar vesicles (GUVs) according to previously published protocols with modifications^5^. The principal lipid used in these experiemnts was soy polar lipids (SPL) or soy polar lipids doped with 20% cholesterol, or a mixture of 70% of POPE and 30% of DOPC. These were supplemented with 0.2 % w/w Rhod-PE (1,2-dioleoyl-sn-glycero-3-phosphoethanolamine-N-(lissamine rhodamine B sulfonyl) (Avanti cat no 810150P) and 0.8 % w/w NBD-PC (1-palmitoyl-2-{6-[(7-nitro-2-1,3-benzoxadiazol-4-yl)amino]hexanoyl}-*sn*-glycero-3-phosphocholine)(Avanti cat no 810130P). 5 mg of lipids were dried with a gentle N_2_ stream on the wall of a glass tube. To completely remove the remaining chloroform, glass tubes were placed in a desiccator for at least two hours at room temperature. Then 1 ml of DR buffer (200 mM KCl, 5 mM HEPES, pH 7.2) was added to the tube. Dried lipids were resuspended by vortexing, then sonicated in a water bath sonicator for 15 minutes to generate small unilamellar vesicles (SUVs). After this, DDM was added to a final concentration of 7 mM to destabilize the SUVs for 30 minutes at room temperature.

Upon destabilization, 4 ml of DR buffer was added to the SUV mixture. The mixture was then aliquoted into 500 µl samples in individual Eppendorf tubes. Proteins were mixed with the lipid mixture of interest at concentrations of (w:w): MscS 1:200; TREK2 1:100; nhTMEM16 1:100; TMEM16F 1:100; TMEM63A 1:50; TMEM63B 1:50; OSCA1.2 1:100. The protein-lipid mixture was incubated with a rotary mixer at room temperature for 1.5 h to allow proteins to insert into the SUVs. After incubation, 50 mg of biobeads (Bio-rad cat no 1523920) were added to each of the Eppendorf tubes and incubated for 1.5 h. Supernatant was collected and 50 mg of fresh biobeads were added to the supernatant for another 1.5 h to fully remove the remaining detergent. The biobeads were discarded after this step and the supernatant was collected for the next step.

### Giant unilamellar vesicle (GUV) formation

For the fluorescence-based scramblase assay, we followed the protocols for gel-assisted dehydration-rehydration as previously described ^89,90^. Glass coverslips with 13 mm diameter were plasma cleaned before use. Low-melting temperature (LMT) agarose was dissolved in water with a concentration of 0.5 % (w/v). 50 µl of melted LMT agarose was evenly spread onto each coverslip and the coverslips were dried at 55 °C for 30 min. After this, 40 µl of protein-lipid mixture from the previous step was carefully spread onto the coverslips and dried in a desiccator for 4 h to overnight at room temperature. Following this, the coverslips were placed into a well of a 24-well plate and 200 µl of DR buffer was added to the well to rehydrate the lipids. The plate was gently agitated at room temperature for 2 h before the GUVs were harvested.

For morphological observations, GUVs were generated with gentle dehydration/rehydration methods. This method yields much less GUVs but does not restrain the liposome morphology in any way. Briefly, 50 µl of protein-lipid mixture was dehydrated on the surface of clean glass slides in a desiccator at room temperature overnight. Following this, 50 µl of DR buffer was added to the dried lipid pellet to rehydrate them into GUVs at room temperature. GUVs were harvested three hours after rehydration.

### Agarose gel-based GUV immobilization

To immobilize GUVs for long term imaging purposes, we follow the protocols previously published with modifications ^91^. Briefly, low melting temperature (LMT) agarose (Sigma-aldrich, cat no A4018), was dissolved in DR buffer to a concentration of 1 % (w/v) at a 99 °C heat block. The LMT agarose was cooled down and maintained on a 37 °C heat block until being used. An equal volume of GUVs were mixed with the melted LMT agarose. For TMEM16F proteoliposomes, 2 mM of EGTA or 1 mM of CaNO_3_ was added to the DR buffer upon rehydration and the LMT agarose gel for immobilization. To maximize diffusion of BSA across the field of view for the scramblase assay below, a 12 well plate with glass-like polymer coverslip bottom (Cellvis) was plasma-cleaned and 80 µl of the agarose-GUV mixture was added to the well to form a thin film. Agarose was allowed to solidify at 4 °C for 20-30 minutes.

### In vitro scramblase assay

Following the steps of immobilization, 80 µl of DR buffer was added to the wells. GUVs were imaged with LSM900 confocal microscope in time series mode at 20 s intervals. After the first three frames, imaging was paused lipid-free BSA (Sigma-Aldrich, cat no A8806) dissolved in DR buffer at a final concentration of 4 mg/ml, doped with 8 µg/ml (as final concentration) of BSA-HiLyte Fluro 647 conjugate (Anaspec, cat no AS-72267), was added to the well to back-extract NBD-PC. 647-BSA was used as a dye to indicate integrity of the GUVs and enable a clear view of BSA diffusion across the well. Another 30 to 40 frames of images were taken until the NBD signal started to plateau. For analysis, the NBD-PC signal was normalized to the Rhodamine-PE signal then normalized to the initial frame. Time courses were plotted with normalized intensity ratio with the endpoint measured at 400 s for quantitative comparison. Alternatively, 1 mM (for soy polar lipids derived GUVs) or 2.5 mM (for GUVs containing 20% of cholesterol) of αCD was added to quench NBD-PC and mechanically stress GUVs. The interval for αCD experiments was 10 seconds.

### Morphological observations of model membranes

Morphological observations of GUVs were carried out with a similar process as described in the previous section. Instead of BSA, 10 µl of 50 mM αCD (final concentration of 5 mM) was added to each well to initiate lipid removal and increase membrane tension and then imaged at intervals of 10 seconds. Recordings lasted for 30 to 40 frames or until GUV lysis. GUVs were classified into ‘broken without tubulation’, ‘broken with tubulation’, ‘shrink without tubulation’, ‘shrink with tubulation’ by visual determination. The ratio of area changed was measured in 2D by normalizing the size of the GUVs at the end of recordings, or one frame immediately before breaking, to their size at the onset of αCD treatment.

### Methods for imaging mTMEM63A mutants

HEK PIEZO1 KO cells were plated in 18-well glass coverslides (81817, Ibidi) coated with fibronectin (10 ug/ml). The next day, they were transfected with the different plasmids (1 mcg/ml of cultured media) using Lipofectamine3000. 24h post transfection, the cultured media was removed and replaced with cultured media containing AnnexinV-FITC (0.3 ug/ml, 640909, BioLegend) and Hoeschst33342 (5 ug/ml, H1399, Invitrogen). The cells were incubated for 10min at 37deg before they were imaged live on a widefield microscope (Eclipe Ti2-E, Nikon) with a 20x objective or on a confocal (LSM900, ZEISS) with a 63x objective.

### Uniaxial stretch

Stretch chambers (SC-0040, STREX) were plasma cleaned at 30 W for 20 s to render them hydrophilic (PDC-002, Harrick Plasma) then immediately coated with gelatin (0.2 % in PBS, G1890, Sigma) and fibronectin (10 µg/ml, F1141, Sigma) at 37 °C overnight. The next day, cells were seeded into PDMS chambers at a density of 120,000 cells/ml and left to attach overnight. The cells were then labelled with AnnexinV-FITC (0.3 µg/ml, 640909, BioLegend) and Hoeschst33342 (5 µg/ml, H1399, Invitrogen) for 10 min at 37 °C. Images were taken before and after stretch using a widefield microscope (Eclipse, Ti2-E, Nikon) and a confocal (LSM900, ZEISS). Stretching was performed uniaxially using a cell stretching device (ST-140-04) at a strain of 10% for 0 – 60 min, in a cell culture incubator. For the cells transfected with OSCA1.1 and 1.2, the cells were transfected 1 day after seeding and were stretched 24 h post transfection.

### Generation of Tmem16f^-/-^ in Piezo1^-/-^ HEK293T cell line

To generate *Tmem16f^-/^*^-^/*Piezo1^-/-^*HEK293T cells, *Piezo1^-/-^* HEK293T cells were transfected with sgRNA targeting the exon 2 of *Tmem16f* and Cas9 protein (RNP complex), in addition to a HDR template containing three stop codons and an EcoRI site, using the Neon electroporation system (Invitrogen). Cells were allowed to rest for 48 h post electroporation then single cell sorted in 96-well plates by plating at a concentration of 0.8 cells/0.1 ml/well. Cells were grown for four to six weeks, and genomic DNA was extracted. Candidate *Tmem16f^-/^*^-^ *Piezo1^-/-^* HEK293T clones were initially validated with PCR with a 20 bp insertion giving a larger PCR product compared to wild type (WT) clones and further validated with Sanger Sequencing and western blotting.

### *In vivo* scramblase assay

*Tmem16f^-/^*^-^/*Piezo1^-/-^* HEK293T cells were transfected with plasmids as indicated. 48 hours post transfection, the cells were digested into single cells and seeded onto a glass bottom 96-well plate at 6,000 cells per well, coated with 0.2% Gelatin and fibronectin (150 nM, Sigma). The cells were allowed to seed for 5 h, and culture media was changed into scramblase buffer (140 mM NaCl, 2 mM CaCl_2_, 2 mM MgCl_2_, 3 mM KCl and 10 mM HEPES adjusted to pH 7.2 using NaOH) containing AnnexinV-568 (0.45 µg/ml, 29010, biotium). αCD or Methyl-β-cyclodextrin (MβCD, sigma) were made into 10x solution to be added into the wells to achieve final concentration as indicated. Real time images were acquired with LSM900 confocal microscope at 1 frame per minute at room temperature. For analysis, the Annexin-V signal from a single cell was normalized to the whole cell area. Normalized intensity in the first frame was designated as the baseline for the following frames. The time courses were plotted with normalized intensity.

### Mechano-resilience assay

Human Oligodendroglioma Cell Line (HOG) (Merck, SCC163) were transfected with control dsiRNA or dsiRNAs targeting TMEM63A (gaccaaagucuacauauu) or TMEM63B (gucaugaccuacaguauc) (IDT) with Lipofectamine RNAiMax (Invitrogen) following manufacturer’s instructions. To maximize knockdown efficiency two treatments with dsiRNA were performed. After 48 h cells were lifted and transfected again with the same amount of transfection reagents and dsiRNA. Cells were then cultured for another 48 h and were digested and seeded onto a glass bottom 96-well plate at 4,500 cells per well, coated with 0.2% gelatin and fibronectin (150 nM, Sigma). The cells were allowed to seed for 5 hours then stained with Hoechst33342 (5 µg/ml, H1399, Invitrogen) for 15 minutes. Culture media was changed into scramblase buffer (140 mM NaCl, 2 mM CaCl_2_, 2 mM MgCl_2_, 3 mM KCl and 10 mM HEPES adjusted to pH 7.2 using NaOH) containing PI (1:100, ab129817) and mounted onto a Nikon imaging system with environmental control (37°C, 5% CO_2_). αCD was added to a final concentration of 25 mM (at 1:1 ratio from a 50 mM stock solution) to rupture the cells. After 30 minutes of αCD addition, PI positive cells and Hoechst positive cells were counted from individual frames and the survived cell proportion was calculated as (N_Hoesct_+ - N_PI+_)/N_Hoechst+_ *100.

