## Supporting Information for "TMEM63 proteins act as mechanically activated cholesterol modulated lipid scramblases contributing to membrane mechano-resilience"

*These authors contributed equally.

#Corresponding authors:

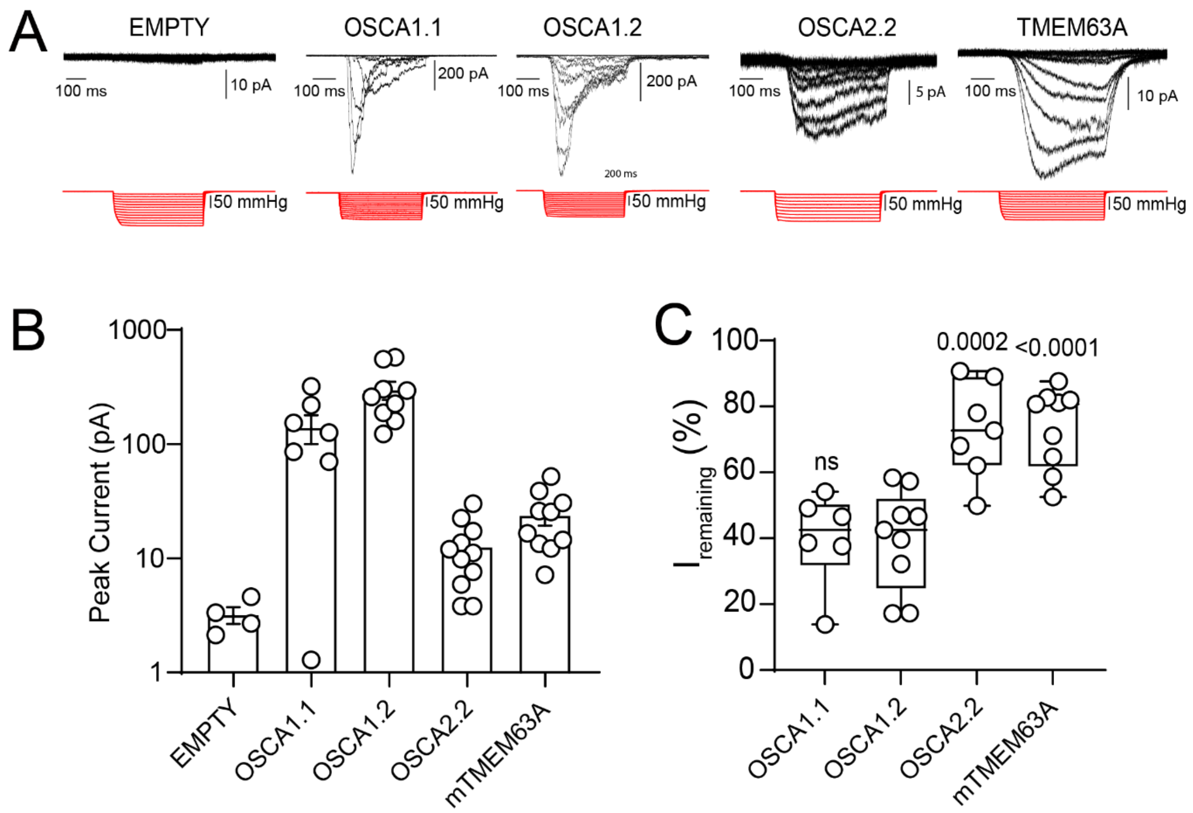


**Figure S1. Comparison of the electrophysiological properties of OSCA1.1/1.2 with OSCA2.2 and TMEM63A.** (A) Representative cell-attached recordings of *Piezo1*$^{-/-}$ HEK293T with an empty vector control, OSCA1.1, OSCA1.2, OSCA2.2 and TMEM63A. All recordings are at a holding potential of -40 mV (B) quantification of the peak current elicited per patch. (C) Quantification of the current remaining ($I_{remaining}$) at the end of the pressure pulse as an indication of the inactivating behavior of OSCA/TMME63 currents. Data shown as max to min box and whiskers plots and p-value determined using one-way ANOVA and Dunnett's post hoc test (ns – non significant).

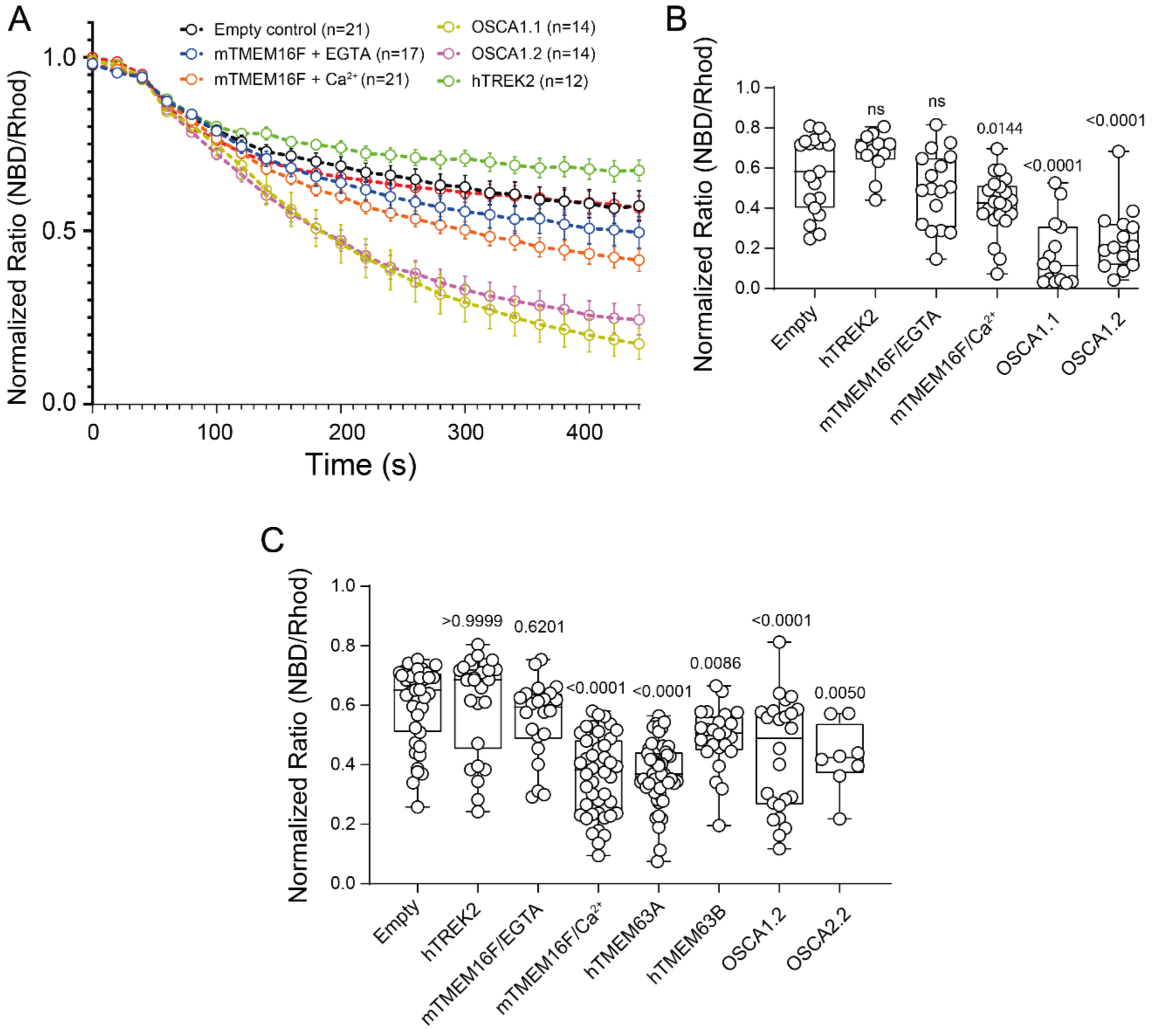


**Figure S2. OSCA1.1 and 1.2 can scramble lipids in GUVs made of PE and PC.**

(A) Time course of changes in normalized ratio of NBD/Rhod fluorescence intensity after BSA addition in empty GUVs (n=19) and those reconstituted with hTREK2 (n=11), mTMEM16F with 2 mM EGTA (n=17), mTMEM16F with 1 mM $Ca^{2+}$ (n=20), OSCA1.1 (n=14) and OSC1.2 (n=14). Error bars represent SEM.

(B) Normalized ratio of NBD/Rhod fluorescence intensity 400 s after BSA addition from the data displayed in panel S2A. Data shown as max to min box and whiskers plots and p-value determined using one-way ANOVA and Dunnett's post hoc test.

(C) Normalized ratio of NBD/Rhod fluorescence intensity 400 s after BSA addition from soy polar lipid GUVs displayed in Figure 1. Data shown as max to min box and whiskers plots and p-value determined using one-way ANOVA and Dunnett's post hoc test.

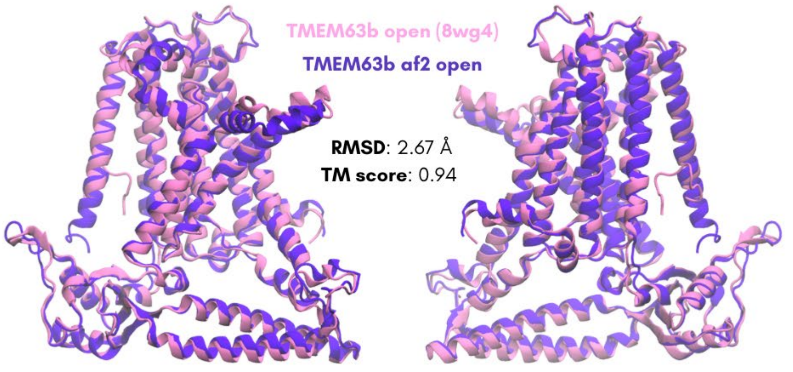


**Figure S3. Structural comparison between a resolved structure of TMEM63b (8wg4) (pink) and AF2 predicted open structure (purple) viewed in two orientations.**

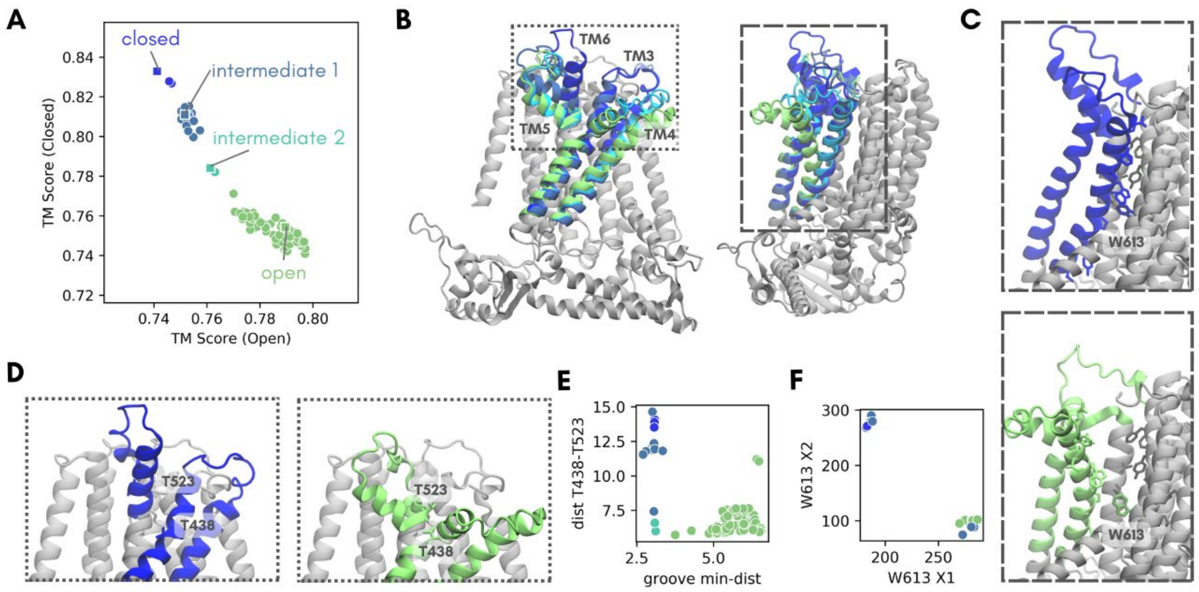


**Figure S4. Structural determinants of TMEM63A functional states**

(A) 100 TMEM63A structures generated by AF2 plotted based on similarity to known open and closed states (as in Fig. 2A). Each structure data point is coloured by its cluster (closed – dark blue, intermediate 1 – ocean blue, intermediate 2 - light blue, open – green), with the central structure of each cluster indicated by a square data point.

(B) Snapshot of four representative TMEM63A structures (closed – dark blue, intermediate 1 – ocean blue, intermediate 2 - light blue, open – green) viewed from the front (left) and side (right).

(C) Differences between groove size and orientation of the W613 side chain in the closed (dark blue, upper panel) and open (green, lower panel) structures as viewed from the side.

(D) Differences in the orientation of TM helices 4-6 between the closed (dark blue) and open (green) structures leads to the formation of an interaction between T438 and T523 in the open state.

(E) Minimum distance between groove-lining residues vs distance between T523 (on TM 6) and T438 (on TM 3) residues on each structure.

(F) Janin plot showing the W613 dihedral angles in each structure

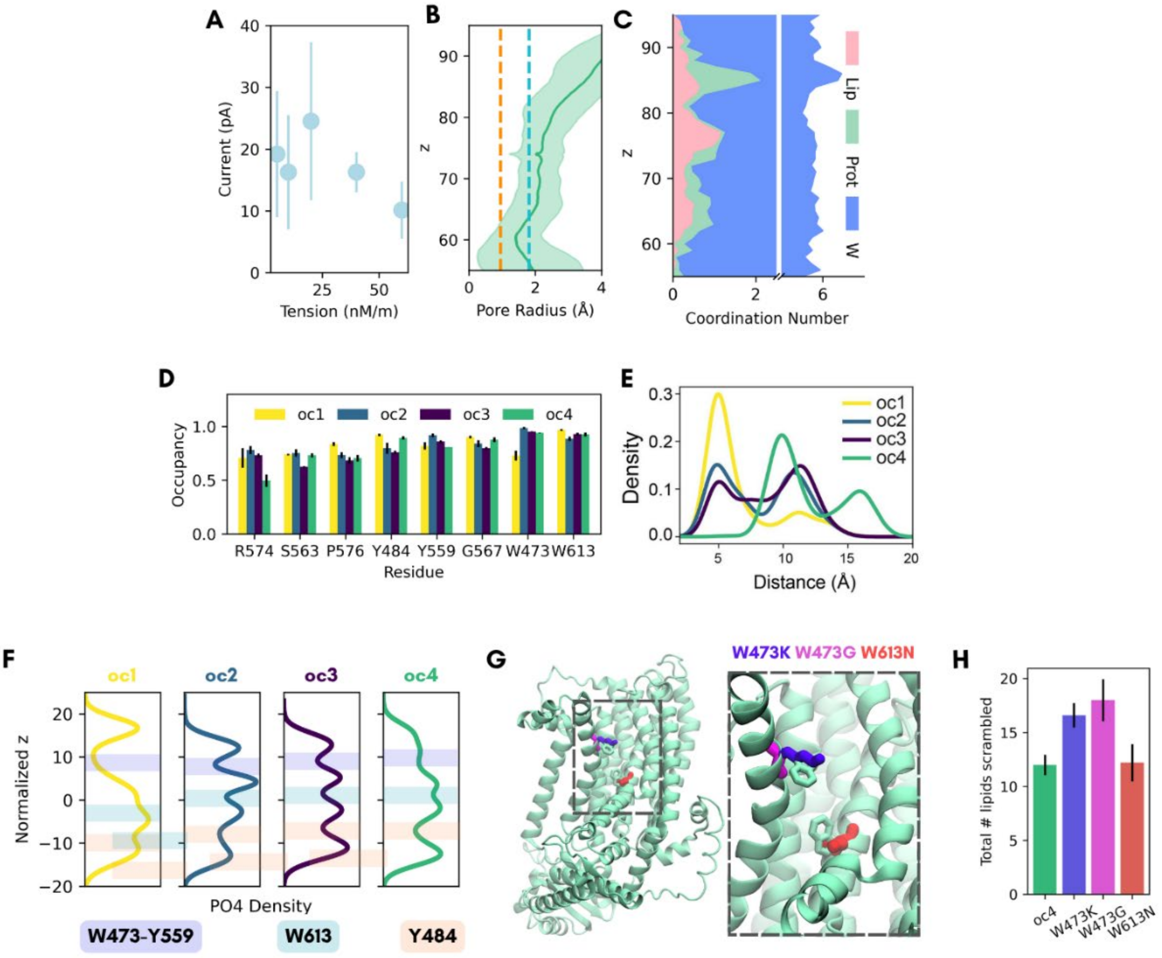


**Figure S5. Structural basis of ion conduction and lipid scrambling in TMEM63A**

(A) Average current through WT TMEM63A at 690 mV across different applied tensions

(B) Pore radius profile for the WT TMEM63A pore at an applied tension of 10 nM/m. The solid green line indicates the mean pore radius, with the coloured bands representing one standard deviation of variation (calculated across every frame of triplicate simulations). The dashed orange line indicates the radius of a dehydrated sodium ion. The dashed blue line indicates the radius of a dehydrated chloride ion.

(C) Coordination number vs pore axis z coordinate, showing the total number of heavy atoms, broken down into those belonging to water, protein and lipid interacting with sodium ions as they permeate the pore.

(D) The proportion of simulation time which groove lining residues are found within 7 Å of translocating lipids. Error bars represent SEM (n=3).

(E) Density distributions of bottleneck distances between side chain beads belonging to bulky residue pairs lining the groove, aggregated across triplicate simulations for each system.

(F) Headgroup (PO4) density at different z positions within the TMEM63A groove throughout triplicate 20 µs simulations for each cluster. The z coordinates were normalised to the average z position of PO4 beads within 2 nm of the protein. Bottleneck locations are indicated by the coloured bars.

(G) TMEM63A oc4 WT structure with key groove residues and mutations made shown in liquorice.

(H) Total number of lipids scrambled in replicate WT and mutant oc4 CGMD simulations of length 40 μs for each system. Error bars represent SEM (n=5).

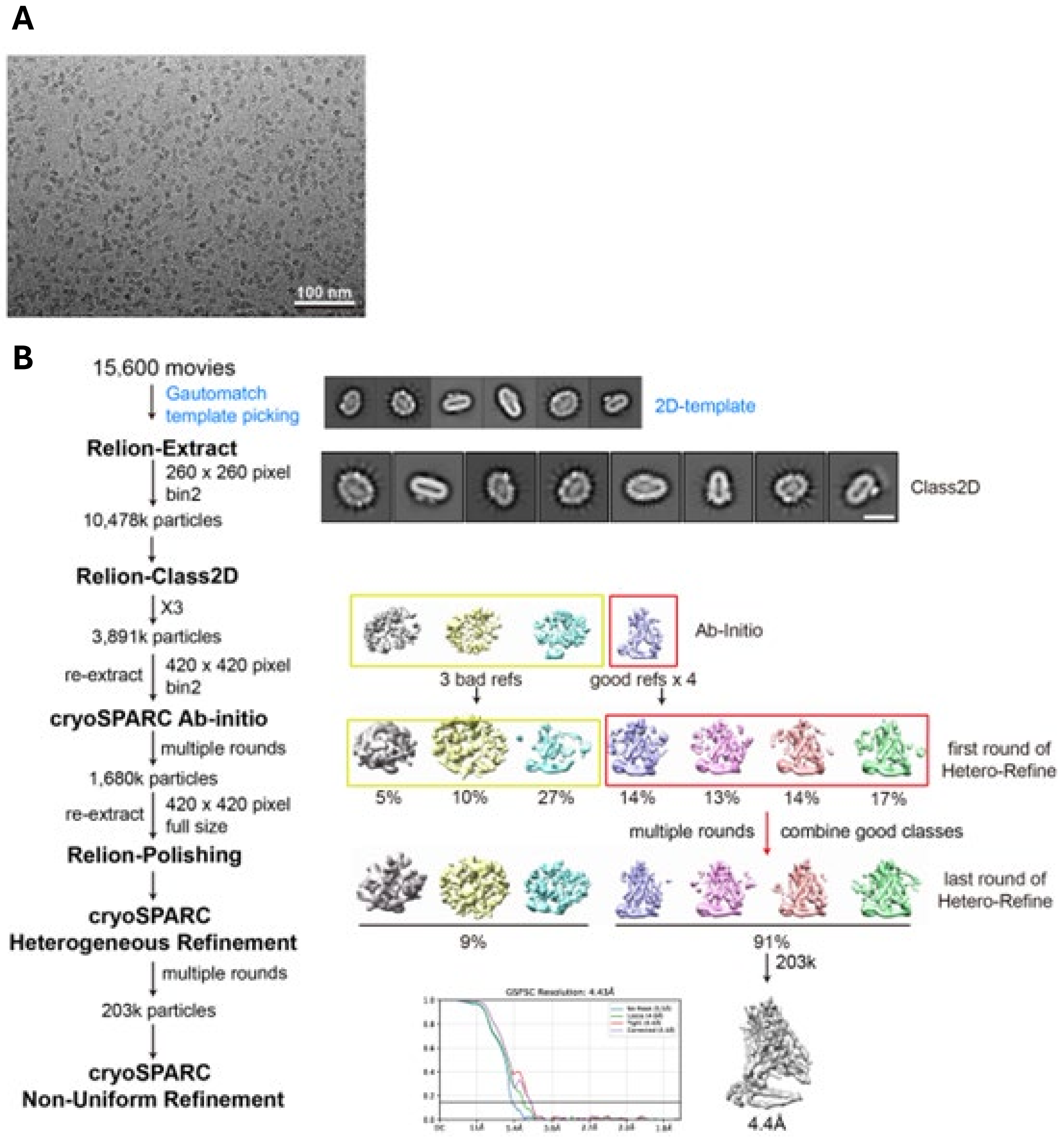


**FigS6. Cryo-EM data processing of TMEM63A-digitonin-cholesterol.**

(A) Representative motion-corrected cryo-EM micrograph of TMEM63A-digitonin-cholesterol. Scale bar, 100 nm.

(B) Image-processing workflow of TMEM63A-digitonin-cholesterol with RELION and cryoSPARC.

Scale bar in the 2D average: 100 nm. Gold-standard FSC curves (FSC = 0.143) are displayed beside the refined maps. See Methods and TableS1 for details.

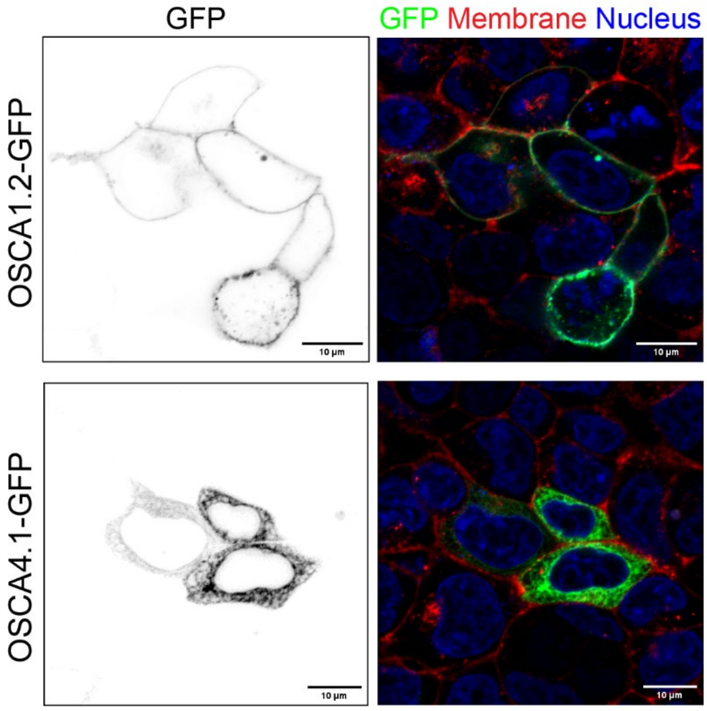


**Figure S7. OSCA4.1 cannot escape intracellular organelles in HEK293T cells.** Transfection of *Piezo1*$^{-/-}$ HEK293T with OSCA1.2-GFP or OSCA4.1-GFP illustrated membrane labelling only in OSCA1.2 expressing cells. The membrane is shown in red and stained with wheat germ agglutinin conjugated to Alexa Fluor 555.

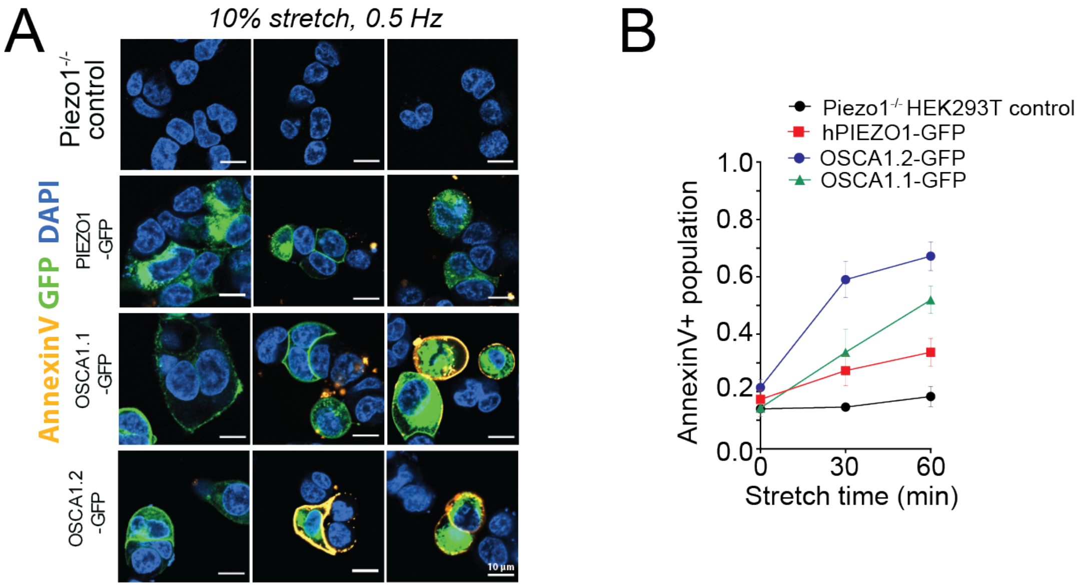


**Figure S8. OSCA scrambling can be driven by mechanical cues *in situ*.**

(A) Confocal images of control *Piezo1*$^{-/-}$ HEK293T cells and the same cells expressing GFP fused versions of OSCA1.1, OSCA1.2 and hPIEZO1 as a negative control after 0, 30 and 60 minutes of cyclic uniaxial stretch at 10% strain and 0.5 Hz. AnnexinV-Alexa568 labelling is represented in gold (scale bar = 10 µm).

(B) Quantification of the normalized amount of transfected cells labelled with annexinV-Alexa568 relative to uniaxial stretch duration. Data points represent mean ± SD (n=3).

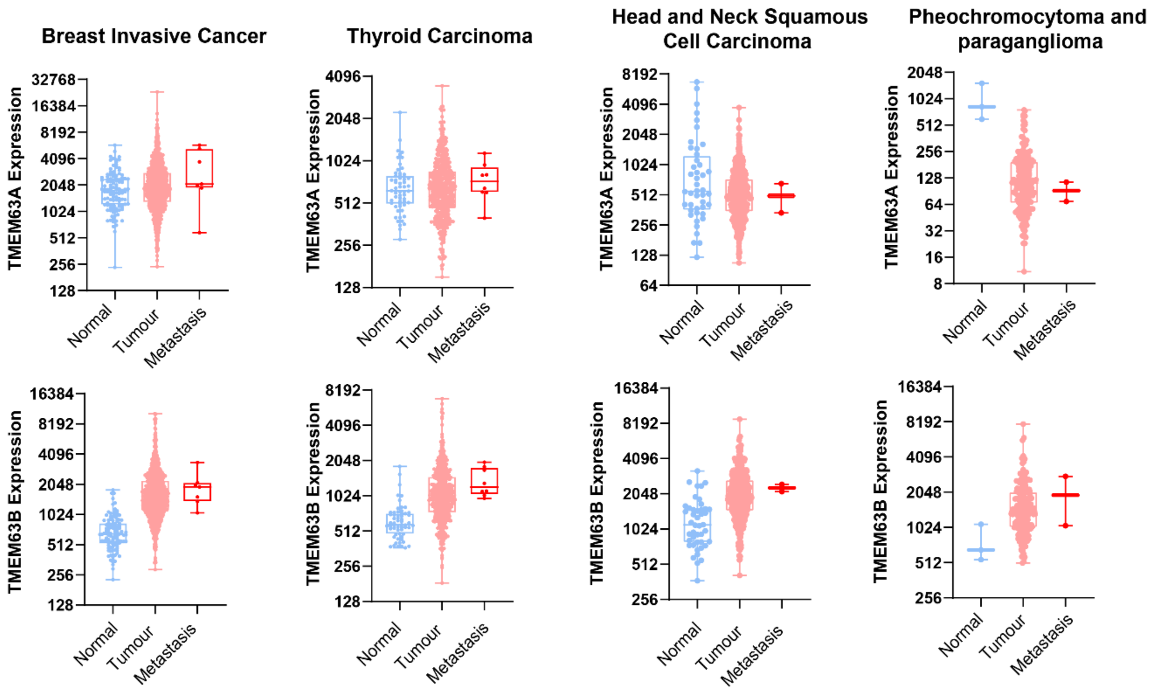


**Figure S9. Correlation of TMEM63A and TMEM63B expression in metastatic cancer cells.** Expression data was extracted from an online repository[1]. The clearest correlation with metastatic cancer is with the plasma membrane localized TMEM63B. Upregulation of TMEM63A in metastatic cancers is only seen in some instances which could be linked to its localization in lysosomes in these cell types. Data shown as max to min box and whiskers plots.

**Table S1. Cryo-EM Data collection, refinement and validation statistics**

| | TMEM63A-dig-cholesterol (EMD-xxxx) (PDB-xxxx) |
|---|---|
| **Data collection and processing** | |
| Magnification | 105,000 |
| Voltage (kV) | 300 |
| Electron exposure (e–/Å$^2$) | 63.8 |
| Defocus range (μm) | -1.4 ~ -2.4 |
| Pixel size (Å) | 1.055 |
| Symmetry imposed | C1 |
| Initial particle images (no.) | 10,478,283 |
| Final particle images (no.) | 202,829 |
| Map resolution (Å) | 4.4 |
| FSC threshold | 0.143 |
| Map resolution range (Å) | 3.0-7.5 |
| **Refinement** | |
| Initial model used (PDB code) | predicted |
| Map sharpening *B* factor (Å$^2$) | -382.4 |
| Model composition | |
| Non-hydrogen atoms | 5152 |
| Protein residues | 626 |
| Ligands | CLR: 2 |
| R.m.s. deviations | |
| Bond lengths (Å) | 0.004 |
| Bond angles (°) | 1.202 |
| Validation | |
| MolProbity score | 2.56 |
| Clash score | 24.71 |
| Poor rotamers (%) | 0 |
| Ramachandran plot | |
| Outliers (%) | 0 |
| Allowed (%) | 15.10 |
| Favored (%) | 84.25 |

**Supplementary Video 1. Lipids are translocated bidirectionally through open OSCA1.2**.

The OSCA1.2 protein backbone is shown using a green surface representation. Lipids in the groove are shown in grey ball and stick representation. The headgroup beads of lipids in the bilayer are shown in transparent grey. Translocating lipids are highlighted in colored ball and stick representation.

**Supplementary Video 2. Lipids are translocated through the AF2 predicted structure of open TMEM63A**

The TMEM63A protein backbone is shown using a green surface representation. Lipids in the groove are shown in grey ball and stick representation. The headgroup beads of lipids in the bilayer are shown in transparent grey. Translocating lipids are highlighted in colored ball and stick representation.
